# Expansion of triplet nucleotide repeats in primates and other vertebrates: an evolutionary perspective

**DOI:** 10.1101/2023.04.05.535742

**Authors:** Saketh Murthy, Rakesh K Mishra

## Abstract

Triplet nucleotide repeat (TNR) expansion has been linked to more than 40 inheritable neurological, neuromuscular and neurodegenerative disorders. Increase in copy number beyond a threshold causes further rapid expansion of the repeats, leading to instability and disease via gain/loss of function, toxic RNA products or chromosome instability. An analysis of these repeat regions across vertebrates shows that these repeats have consistently either arisen late or have increased in copy number in vertebrates, most significantly in primates and particularly in humans. Many of the known diseases have neurological basis, suggests positive selection of these repeats for neuronal function. Late occurrence of the diseases implicates a lack of negative selection. This evolutionary trade-off, a higher neuronal capability at the cost of disease susceptibility, is further supported by the observation that most of the genes associated with TNR expansion diseases have neuronal function.

## INTRODUCTION

TNR expansion is a kind of mutation in which repeats of three nucleotides increase in copy numbers. Once the copy number crosses a threshold, the repeat tract becomes unstable causing further expansion resulting in disorders referred to as triplet nucleotide repeat disorders (TNR disorders). Repeat instability is linked to more than 40 inheritable neurological, neurodegenerative and neuromuscular disorders^[1]^. The repeat mutation process is dynamic, with products that continue to mutate within tissues and across generations. TNR disorders cause disease via gain/loss of function, toxic RNA products or chromosome instability^[2]^.

Triplet expansion can occur during DNA replication, DNA repair and meiotic recombination. Due to the repetitive nature of the DNA sequences in these repeat regions, ‘loop out’ structures may form during DNA replication while maintaining complementary base pairing between the parent strand and the daughter strand being synthesized, with repeats expanding or contracting based on whether the loop is formed on the daughter or parent strand, respectively^[3]^. DNA repair processes such as homologous recombination, non-homologous end joining, mismatch repair or base excision repair involve a DNA synthesis step where strand slippage can lead to TNR expansion^[4]^. Unequal homologous exchange during meiotic or mitotic recombination can also cause repeat expansion. Some other mechanisms for expansion and reduction have been proposed involving interaction of RNA and DNA molecules^[5]^. Variability in copy number of repeats within a tissue (mosaicism) indicates instability has occurred but not at what stage. However, age-dependent instability, with repeats accumulating in post-mitotic tissues (neurons), implicates genome-maintenance repair as a cause of repeat expansion^[1]^.

In terms of location, expanding TNRs can be found in both coding and non-coding regions of disease-causing genes. In the coding regions, mostly CAG and CGN repeats are present, leading to polyglutamine and polyalanine tracts respectively^[6]^. At the 5’ UTR, CGG and CAG repeats are present, while at the 3’ UTR, CTG repeats are found. Various triplet repeats are also found in the intronic and promoter regions^[6]^. Of the coding-region repeats, there are two classes. Polyglutamine repeat expansions cause a variety of neurological disorders, and polyalanine repeat expansions are implicated in developmental disorders. The non-coding region repeats also cause neurological and developmental disorders^[1]^.

In TNR expansion, once the repeat number crosses a certain threshold (typically 30-40 repeats), the repeats start to rapidly expand causing longer and longer expansions in future generations. But if the number of repeats is below the threshold, it remains relatively stable. The number of triplet repeats can typically increase to around 100 in coding regions and up to thousands in non-coding regions^[7]^. This difference is due to over-expression of glutamine and alanine, which is selected against due to cell toxicity^[8]^. The number of trinucleotide repeats appear to predict the progression, severity, and age of onset of several of the TNR disorders^[1]^. It has been shown that there is a clear inverse relationship between the length of the repeats in parents and the age of disease onset in children, though high variability exists among patients^[9,10]^. This phenomenon of genetic anticipation is a hallmark of TNR disorders.

Triplet nucleotide disorders are uniquely prevalent in humans. There is a lack of evidence of TNR disorders in other organisms. Most of the diseases have a neurological basis and are inheritable late-onset diseases. These intriguing facts led us to study the presence of these repeats in other vertebrates to investigate the evolutionary significance/evolutionary basis of these disorders. In this study, we investigate 18 known TNR disorders that occur as a result of triplet nucleotide expansions in the exonic regions of protein-coding genes, and 7 known TNR disorders that occur as a result of expansion in non-exonic regions, and analyze the presence and length of these repeats across vertebrates.

## METHODS

The analysis was performed on 25 genes with repeats involved in triplet nucleotide repeat expansion related diseases in humans (*Table 1*). 18 of these genes have repeat expansion in the exons, and the 7 others have in the non-coding regions. The analysis was done using two separate strategies.

**Table 1.**
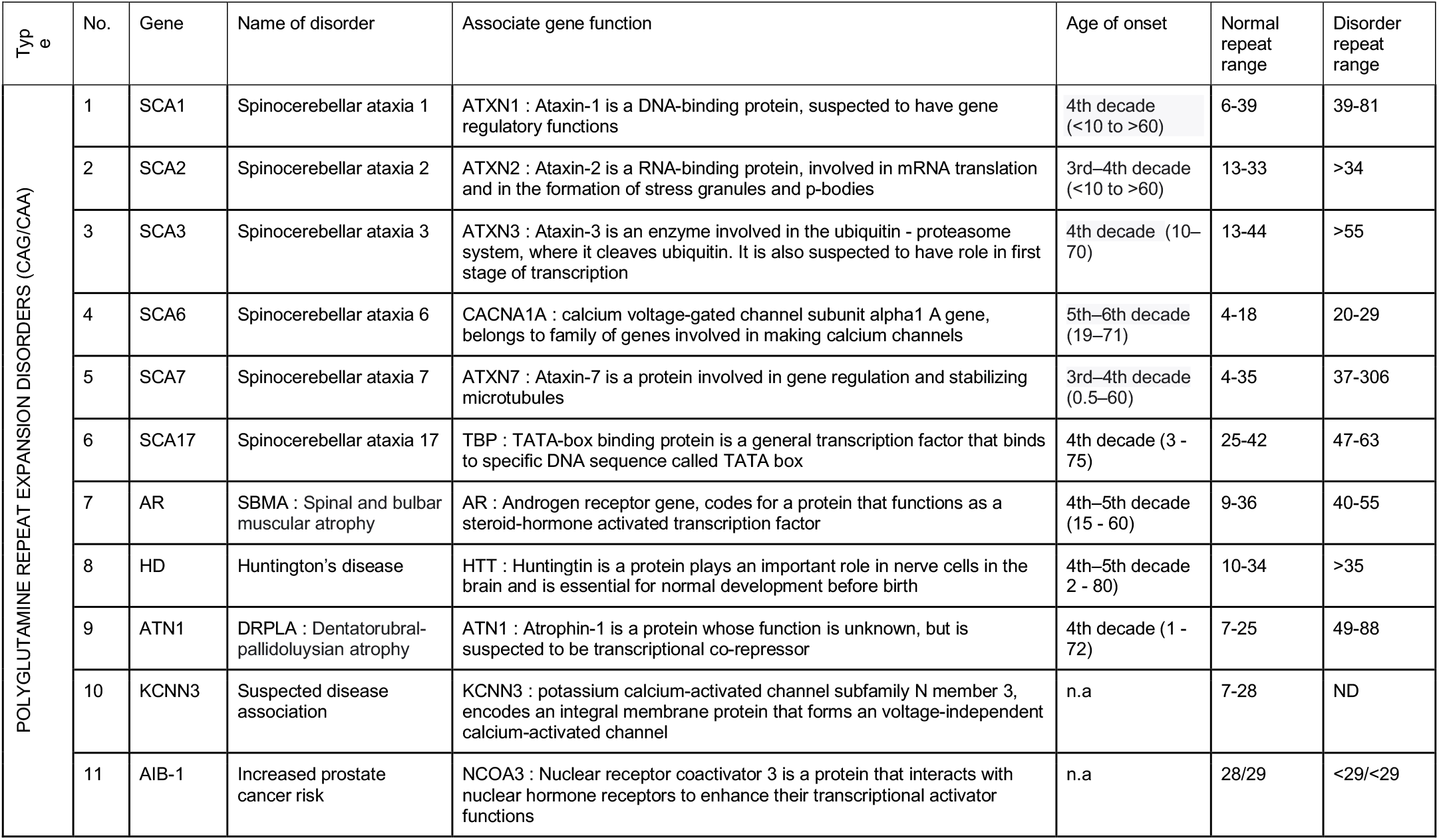
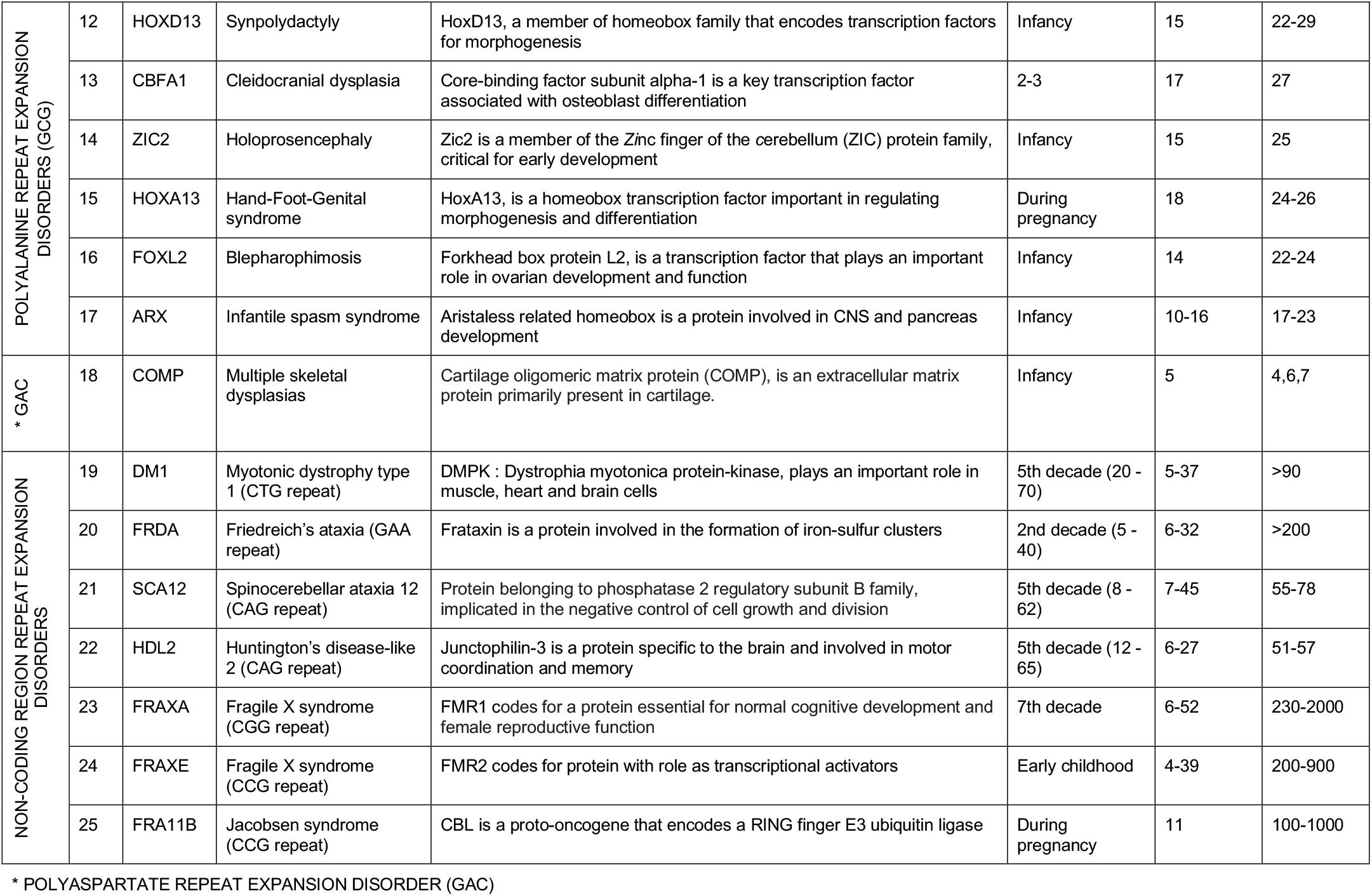
List of coding-region and non-coding region triplet nucleotide expansion related diseases in humans

### BLASTn to identify orthologs

In this strategy we use BLASTn to identify orthologs across a select number of organisms. 36 model organisms were chosen for the analysis spanning vertebrates (*Table 2*). These organisms were chosen based on the presence of reference quality genome assemblies on the NCBI database and spanning the major vertebrate families with a focus on primates and mammals. First, the human protein sequences of the 18 genes were downloaded. Next BLASTn was performed using the repeat regions and 150 bases on either end, to pick out homologs in the 36 organisms, using default parameters on the NCBI website^[14]^. All sequences were downloaded from the NCBI website as of 8 April, 2021. Two criteria were applied to determine whether a homolog was used in the analysis : 1) A sequence match of high significance (e-value < 10^-5) and 2) the picked homolog sequence should belong to a reference assembly of the organism. Some of the homologs were based on prediction methods (rather than experimentally validated), but were used only if the base assembly was of reference quality. These criteria, and the absence of blast hits in some cases, meant the absence of protein homolog data for some of the organisms. I used the multiple alignment tool available on the NCBI website, and used the match to locate the repeat regions and count the repeat lengths in each homolog. For each of the homologs, the maximum length of uninterrupted tandem repeats were counted and recorded. I have used lengths of uninterrupted repeats as previous studies have shown that disease phenotypes are associated with pure, uninterrupted tracts.^[11, 12]^

**Table 2.** List of organisms chosen for the analysis

|  |  |  |  |  |
| --- | --- | --- | --- | --- |
| Vertebrates | Mammals | Primates | Apes | Human (*Homo sapiens*), Chimpanzee (*Pan troglodytes*), Orangutan (*Pongo abelii*), Gorilla (*Gorilla gorilla*), Gibbon (*Nomascus leucogenys*) |
|  |  |  | Monkeys | Rhesus Macaque (*Macaca mullata*), Squirrel Monkey (*Saimiri boliviensis*) |
|  |  | Rodents |  | House mouse (*Mus musculus*), Rat (*Rattus norvegicus*), Guinea Pig (*Cavia porcellus*), Squirrel (*Ictidomys tridecemlineatus*), Rabbit (*Oryctolagus cuniculus*) |
|  |  | Carnivores |  | Dog (*Canis lupus*), Ferret (*Mustela putorius*), Cat (*Felis catus*), Giant Panda (*Ailuropoda melanoleuca*) |
|  |  | Other Mammals |  | Pig (*Sus scrofa*), Cow (*Bos taurus*), Horse (*Equus caballus*), Shrew (*Sorex araneus*), Megabat (*Pteropus vampyrus*), Chinese Pangolin (*Manis pentadactyla*), Dolphin (*Tursiops truncatus*), Elephant (*Loxodonta africana*) |
|  | Non Mammals | Birds |  | Zebra Finch (*Taeniopygia guttata*), Chicken (*Gallus gallus*), Turkey (*Meleagris gallapavo*), Golden Eagle (*Aquila chrysaetos*) |
|  |  | Reptiles |  | Lizard (*Anolis carolinensis*), Garter Snake (*Thamnophis sirtalis*) |
|  |  | Amphibians |  | High Himalayan Frog (*Nanorana parkeri*), Western Clawed Frog (*Xenopus Tropicalis*) |
|  |  | Fish |  | Zebra Fish (*Danio rerio*), Fugu (*Takifugu rubripes*), Medaka (*Oryzias latipes*), Atlantic Cod (*Gadus morhua*) |

### OrthoDB to identify orthologs

In this strategy we use OrthoDB^[15]^ to access the vertebrate orthologs. OrthoDB was used to access all the vertebrate orthologs of each of 18 genes with repeats in the exonic regions, and their amino acid sequences were downloaded on Dec 20, 2022. Uninterrupted repeats of minimum 3 amino acid length were identified in all the orthologs. When there is 2 or less amino acid repeat stretch, it is taken as 0 for further averaging and plotting.

## RESULTS

The analysis was done for the 18 known TNR diseases involving expansion of repeats in exonic regions of genes and 7 known TNR diseases involving expansion of repeats in non-coding regions of genes. For the BLASTn-led approach, 36 vertebrates were chosen and the length of the repeats were compared across these vertebrates. For the OrthoDB-led approach, all available vertebrate orthologs were taken, and the repeat lengths compared.

The lengths of uninterrupted tandem repeats for each organism and protein using the BLASTn approach are recorded and plotted in (*Supp. files 1 and 2*). The length of tandem repeats using the OrthoDB approach is plotted in (*Supp files 3 and 4*). For some of the protein-organism combinations, BLASTn did not give a hit: this could be because the protein is either absent or has diverged widely in these organisms. Such combinations have been marked with an ‘x’ and ‘?’. All the organisms were split into 5 classes (human, other apes, other primates, other mammals, and non-mammals) and the average repeat length for each class was calculated and plotted along with that of humans below.

### Polyglutamine repeats

These are disorders caused by expansion of glutamine and lead to neurological disorders. The repeat length variation across vertebrates for the 11 known polyglutamine repeat expansion related disease genes were analyzed in this study, *Figure 1*. An analysis of the plots throws up some striking observations. All the genes are present and functioning across vertebrates, regardless of the presence and length of the repeats. When comparing the non-mammals to the mammals; the repeats are either completely absent or when present, the repeat lengths are very short in the nonmammals. When comparing primates to other mammals, we see a trend of significant expansion of repeats in the primates. Among the primates, humans have the highest repeat length in 8 of the 11 genes. Overall, there is a clearly visible trend of absence or highly reduced presence of the repeats in non-mammals, some expansion in mammals, and a marked increase in the primates, with humans and chimps having the highest levels of expansion.

**Figure 1.**
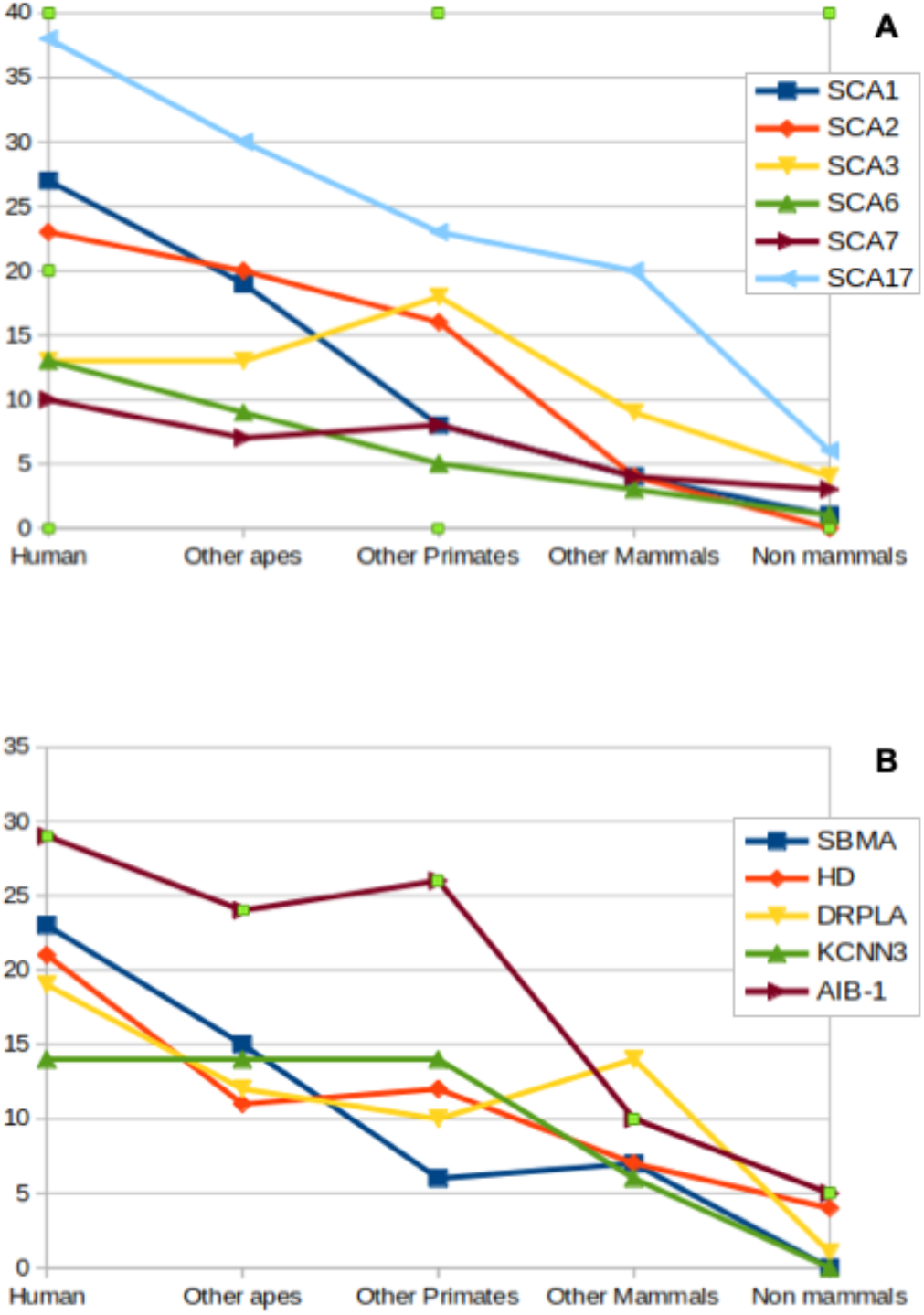
Polyglutamine repeat lengths in TNR associated genes across vertebrates: A. Spinocerebellar ataxias, B. Other disorders. The repeats are either absent or present with very low copy numbers in non-mammals. In primates, the repeats are significantly expanded. The highest copy numbers are found in humans and chimps for every repeat.

### Polyalanine/Polyaspartate repeats

These are disorders caused by expansion of alanine and aspartate leading to developmental defects. The repeat length variation across vertebrates for the 6 known polyalanine repeat expansion related disease genes and the 1 known polyaspartate repeat expansion related disease gene (COMP) were analyzed in this study, *Figure 2*. All genes have homologs across the vertebrates, irrespective of presence or length of repeats. As in the case of the polyglutamine genes, all the polyalanine genes have either no repeats or highly reduced repeat lengths in non-mammals. The repeats are expanded in the mammals and all the mammals have highly conserved repeat lengths with almost no variation amongst them. Many of these genes play important developmental roles, which may explain the tight regulation of the number of repeats. In the case of COMP, the conserved repeat lengths extend to the non-mammals as well.

**Figure 2.**
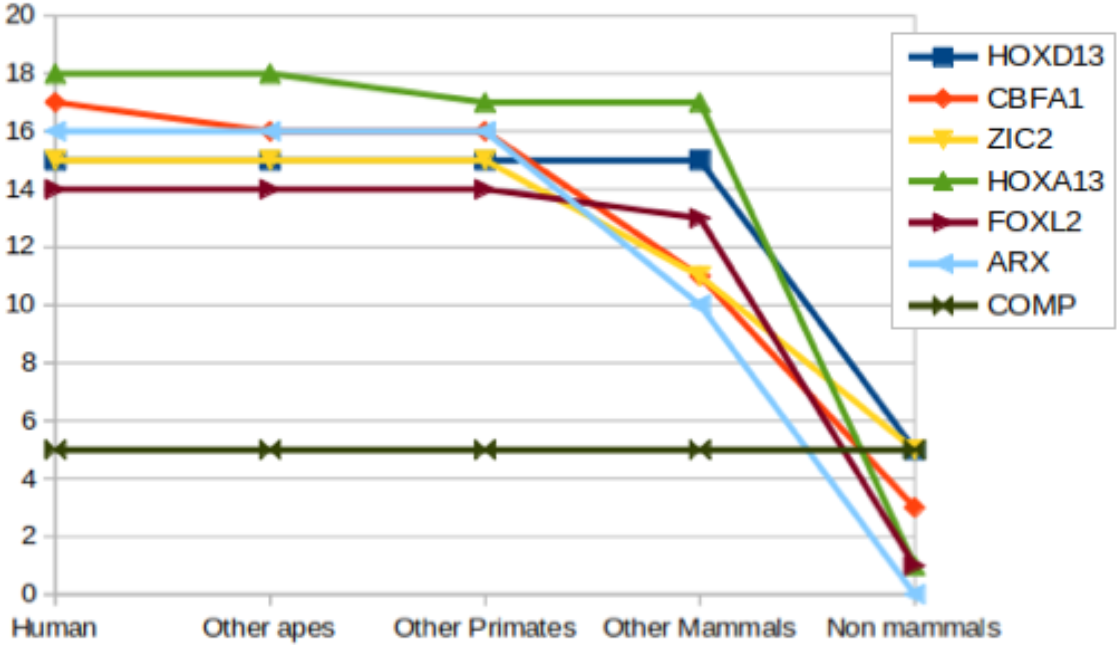
Repeat lengths of the polyalanine/polyaspartate repeat tracts in disease-causing genes. The repeats are either absent or present with very low copy numbers in non-mammals. In mammals, the repeats are present and the repeat lengths are highly conserved.

### Non-coding region repeats

These are disorders caused by expansion of various repeat motifs in the untranslated regions of the genes. They are neurological and developmental disorders. The repeat length variation across vertebrates for the 7 known non-coding region repeat expansion related genes were analyzed in this study, *Figure 3*. All genes have homologs across the vertebrates, irrespective of presence or length of repeats. In the non-coding region repeats also, we see a similar pattern to the coding region repeats. There is a clearly visible trend of absence or highly reduced presence of the repeats in non-mammals, some expansion in mammals, and a marked increase in the primates.

**Figure 3:**
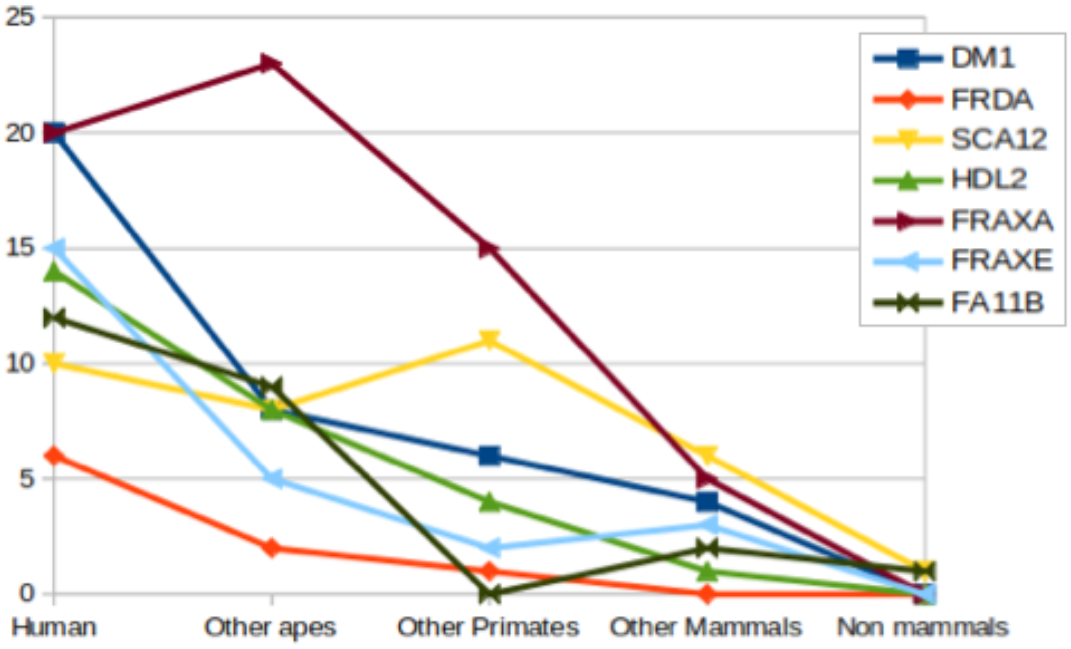
Repeat lengths of non-coding region repeat tracts in disease-causing genes across vertebrates. The repeats are either absent or present with very low copy numbers in non-mammals. In primates, the repeats are significantly expanded.

## DISCUSSION

With the evolution complexity, there are a number of novel mechanisms present in higher eukaryotes and vertebrates, which are absent in invertebrates and more ancient lineages. The way the chromosome is packed and interpreted has evolved significantly over evolutionary time. Gene regulatory networks, elements of gene regulation and control of gene expression, post-translational modifications, epigenetic modifications, non-coding RNA elements, chromosome packing and arrangement, have all seen significant expansion and diversification in higher eukaryotes, especially vertebrates.

Repeat expansion-related disorders are a class of disorders dependent on expansion of repeat regions beyond a certain threshold. The 18 known genes involved in disease are all conserved across vertebrates. Polyglutamine repeat expansion disorders have a neurological basis, and the Polyalanine repeat expansion disorders are involved in embryonic development. Repeat disorders are unique to humans, and many disorders have a strong neurological basis. This begs the question - what causes these disorders to evolve and be maintained specifically in humans? Is there any significance to their neurological basis?

When we compare the lengths of the repeats across the vertebrates, we see a few striking features. First, for all the genes, the repeats either don’t exist or exist in very low length in the non-mammals. Second, the repeat length increases in mammals and is found to be the highest in primates and in humans. A study of the prevalence of repeats in proteins has shown that a higher fraction of proteins have repeats in eukaryotes, especially vertebrates. But a study of repeat lengths across proteins shows that amongst all repeat classes, only GAA repeats have expanded in mammals, and CAG repeats are not expanded in any particular lineage.^[13]^

Our analysis shows that these repeat regions within exonic regions of disease-causing genes have expanded in mammals and especially primates, across all the genes studied. This suggests an evolutionary role for these repeat regions in these proteins. As neurological function has expanded and evolved, it appears that repeat regions have evolved for certain biological functions in neurons. The tight control of the number of repeats in polyalanine tracts also points to a positive functional role of these repeats in the proteins.

Repeats are seen to expand close to and beyond the threshold of disease in humans only. The high prevalence and occurrence of various old-age diseases in humans implicate the lack of a strong negative selection pressure, with the diseases occurring long after reproductive age. TNR diseases also belong to this class of diseases, with early onset of diseases only occurring after a number of generations of expansion. The inheritance of these diseases, the phenomenon of genetic anticipation, and the old-age onset of many of the TNR diseases suggests that these repeats have while evolved under the positive selection pressure, probably related to the distinct neuronal capability of primates, and in particular humans, they are maintained in the population due to a lack of negative selection.

We also explored the regions/tissues for the expression of these genes to further correlate with their function. Most these gene have ubiquitous expression throughout, with a few notable exceptions. For example POLYGLU repeat genes are expressed throughout all tissues but for CACNA1A and KCNN3 that have high expression in the brain. AR had low expression in brain and high in liver, endometrium, ovary, and prostate. Other TNR genes except the ones below were expressed throughout tissues. HOXD13 and HOXA13 are expressed only in colon, prostate, urinary bladder, while ZIC2 is expressed high only in brain, and testes. HDL2 is expressed high in brain and ARX is expressed in ovary, brain, stomach, and testes. COMP has broad expression across 11 tissues, but low in brain. While these observations do indicate that the function of TNR expansion related disease genes are largely associated with neuronal tissues, their additional functions may not be ruled out. While these observations do indicate that the function of TNR expansion related disease genes are largely associated with neuronal tissues, their additional functions may not be ruled out.

In this study, we compared uninterrupted, pure stretches of codon repeats. Various studies have found that interrupts increase stability of the alleles. It would be interesting to see in which lineages and where interruptions arise, and if they have any role in stability or expansion of repeat stretches in the context of the function of the associated gene.

In summary, we have two class of TNR expansion related gene, i. Polyglu genes and most of the non-coding region genes, show neurological diseases and old-age onset, and we hypothesize that neuronal function as positive selection and old age disease as lack of negative selection; ii. The polyala genes and some of the non-coding region genes, show developmental disorders and early-age onset. We still posit a positive selection, as repeats are absent in lower vertebrates like fishes. We also see tight control of repeat numbers in mammals, which we attribute to them being developmentally crucial. Taken together, these observations of the repeat lengths of triplet nucleotides in disease causing genes across vertebrates, point to the dual role of positive selection of repeat tracts in these genes for neuronal activity, and a lack of negative selection of the expanded repeats in these genes largely due to post-reproductive stage pathology of these diseases. It also provides an interesting example of evolutionary trade-off - intelligence, at the cost of old age disease susceptibility, seems to have prevailed.

## ACKNOWLEDGMENTS

Authors acknowledge Akshay Avvaru for comments and technical help. RKM lab research is supported by CSIR, New Delhi.

