## Supplementary file 2 for "Expansion of triplet nucleotide repeats in primates and other vertebrates: an evolutionary perspective"

### POLYGLUTAMINE TNR RELATED  DISEASES

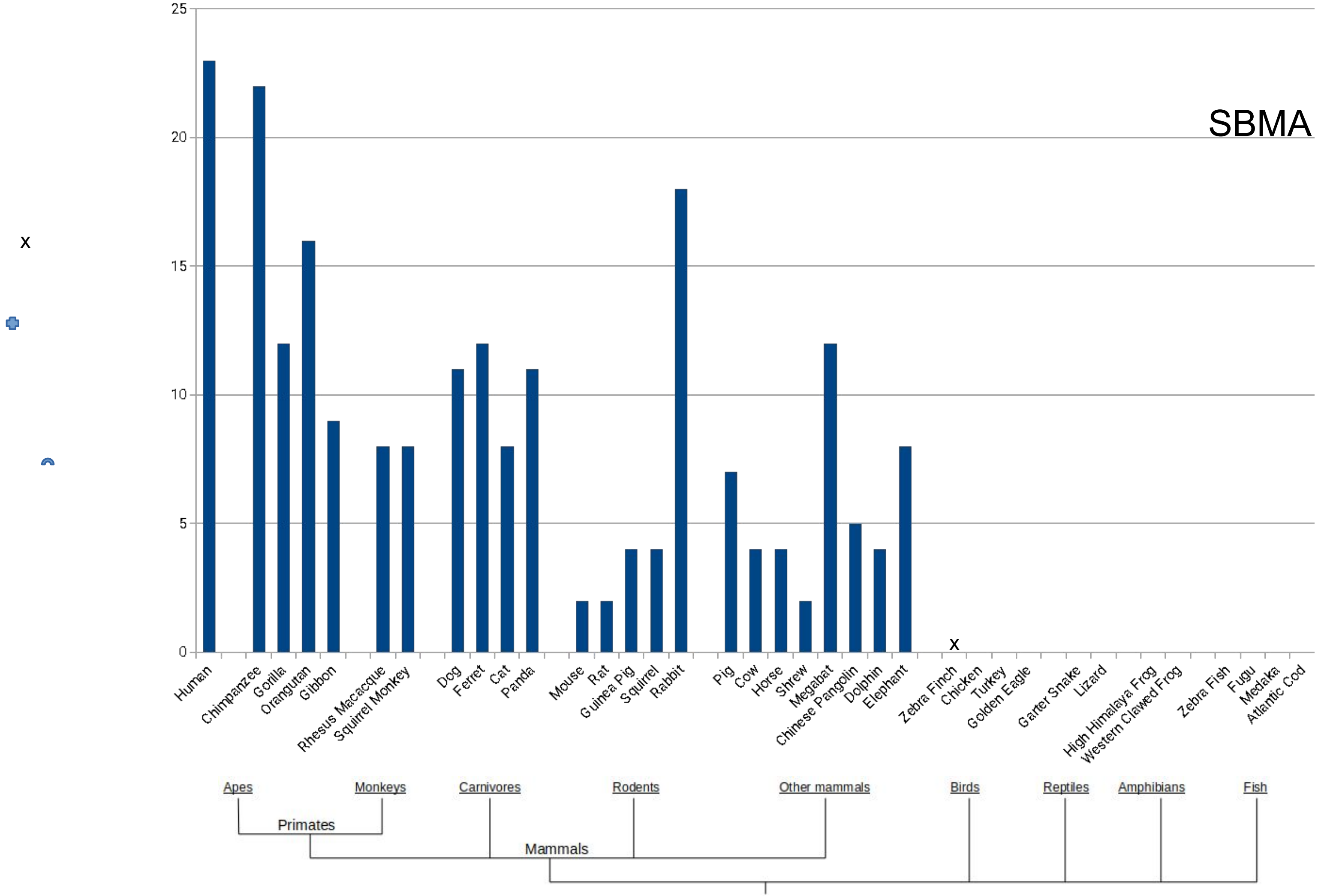

SBMA
25
20
15
10
5
0
x
x
Human
Chimpanzee
Gorilla
Orangutan
Gibbon
Rhesus Macacque
Squirrel Monkey
Dog
Ferret
Cat
Panda
Mouse
Rat
Guinea Pig
Squirrel
Rabbit
Pig
Cow
Horse
Shrew
Megabat
Chinese Pangolin
Dolphin
Elephant
Zebra Finch
Chicken
Turkey
Golden Eagle
Garter Snake
Lizard
High Himalaya Frog
Western Clawed Frog
Zebra Fish
Fugu
Medaka
Atlantic Cod
Apes
Monkeys
Carnivores
Rodents
Other mammals
Birds
Reptiles
Amphibians
Fish
Primates
Mammals

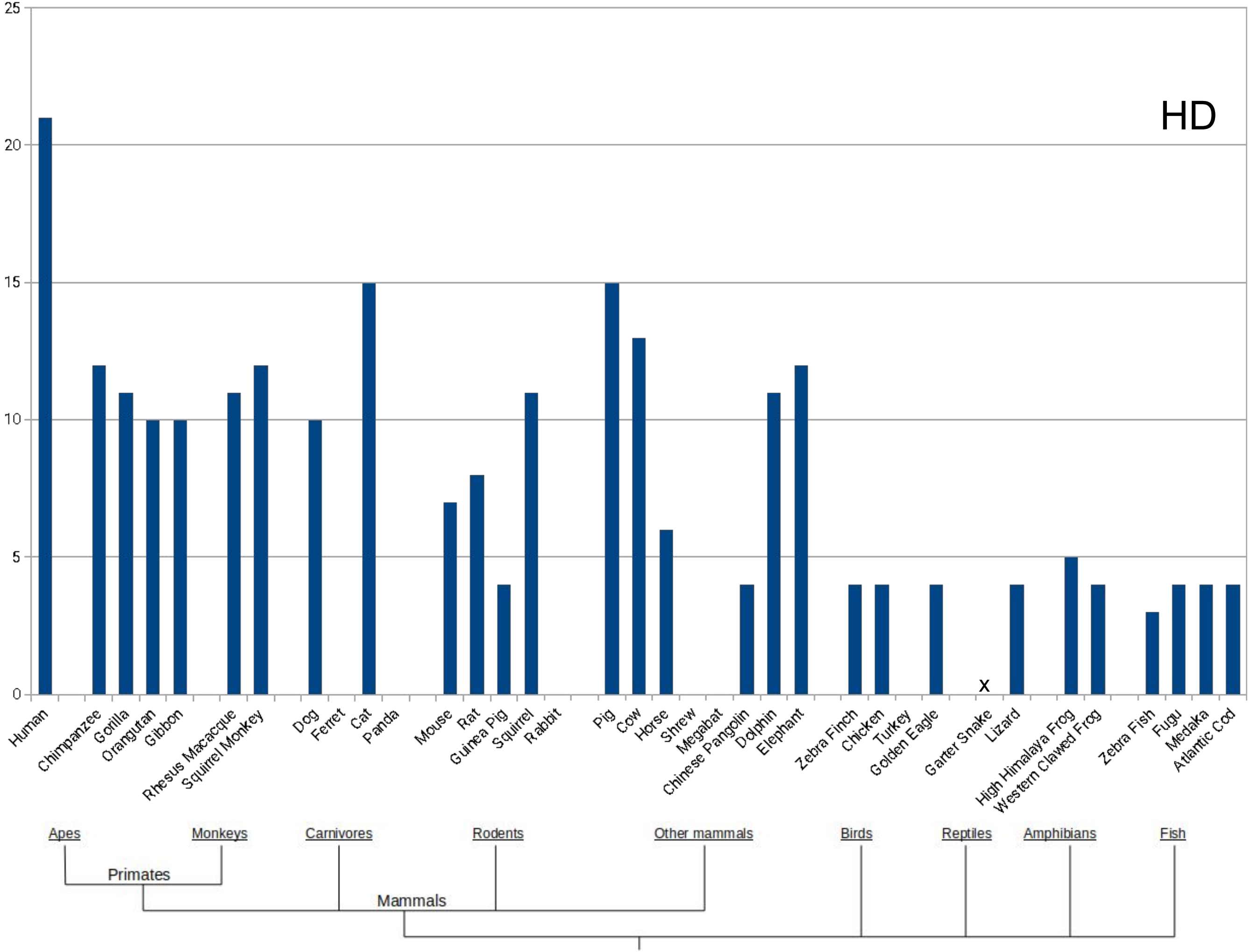

HD
25
20
15
10
5
0
x
Human
Chimpanzee
Gorilla
Orangutan
Gibbon
Rhesus Macacque
Squirrel Monkey
Dog
Ferret
Cat
Panda
Mouse
Rat
Guinea Pig
Squirrel
Rabbit
Pig
Cow
Horse
Shrew
Megabat
Chinese Pangolin
Dolphin
Elephant
Zebra Finch
Chicken
Turkey
Golden Eagle
Garter Snake
Lizard
High Himalaya Frog
Western Clawed Frog
Zebra Fish
Fugu
Medaka
Atlantic Cod
Apes
Monkeys
Carnivores
Rodents
Other mammals
Birds
Reptiles
Amphibians
Fish
Primates
Mammals

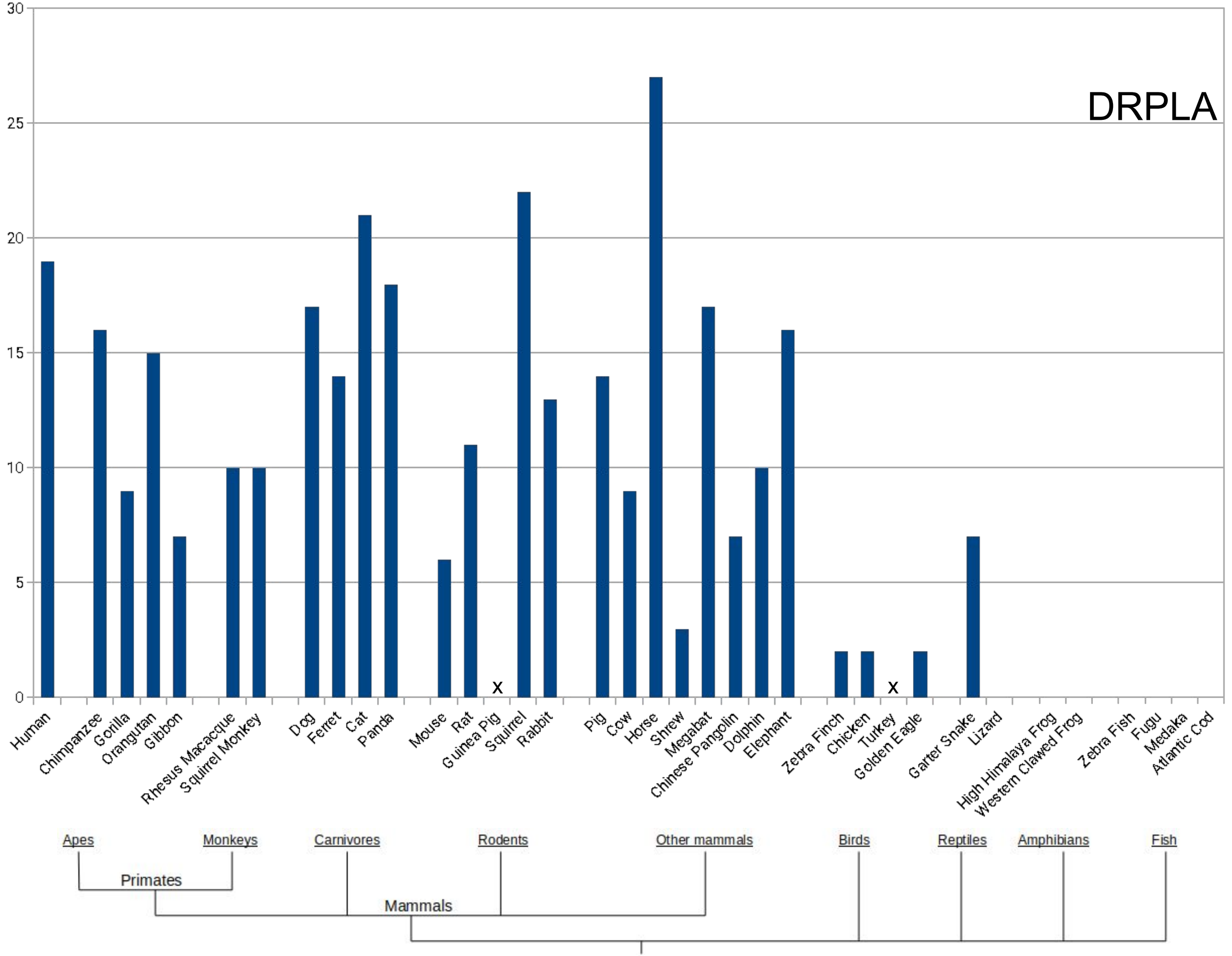
DRPLA
30
25
20
15
10
5
0
Human
Chimpanzee
Gorilla
Orangutan
Gibbon
Rhesus Macacque
Squirrel Monkey
Dog
Ferret
Cat
Panda
Mouse
Rat
Guinea Pig
Squirrel
Rabbit
Pig
Cow
Horse
Shrew
Megabat
Chinese Pangolin
Dolphin
Elephant
Zebra Finch
Chicken
Turkey
Golden Eagle
Garter Snake
Lizard
High Himalaya Frog
Western Clawed Frog
Zebra Fish
Fugu
Medaka
Atlantic Cod
x
x
Apes
Monkeys
Primates
Carnivores
Rodents
Other mammals
Mammals
Birds
Reptiles
Amphibians
Fish

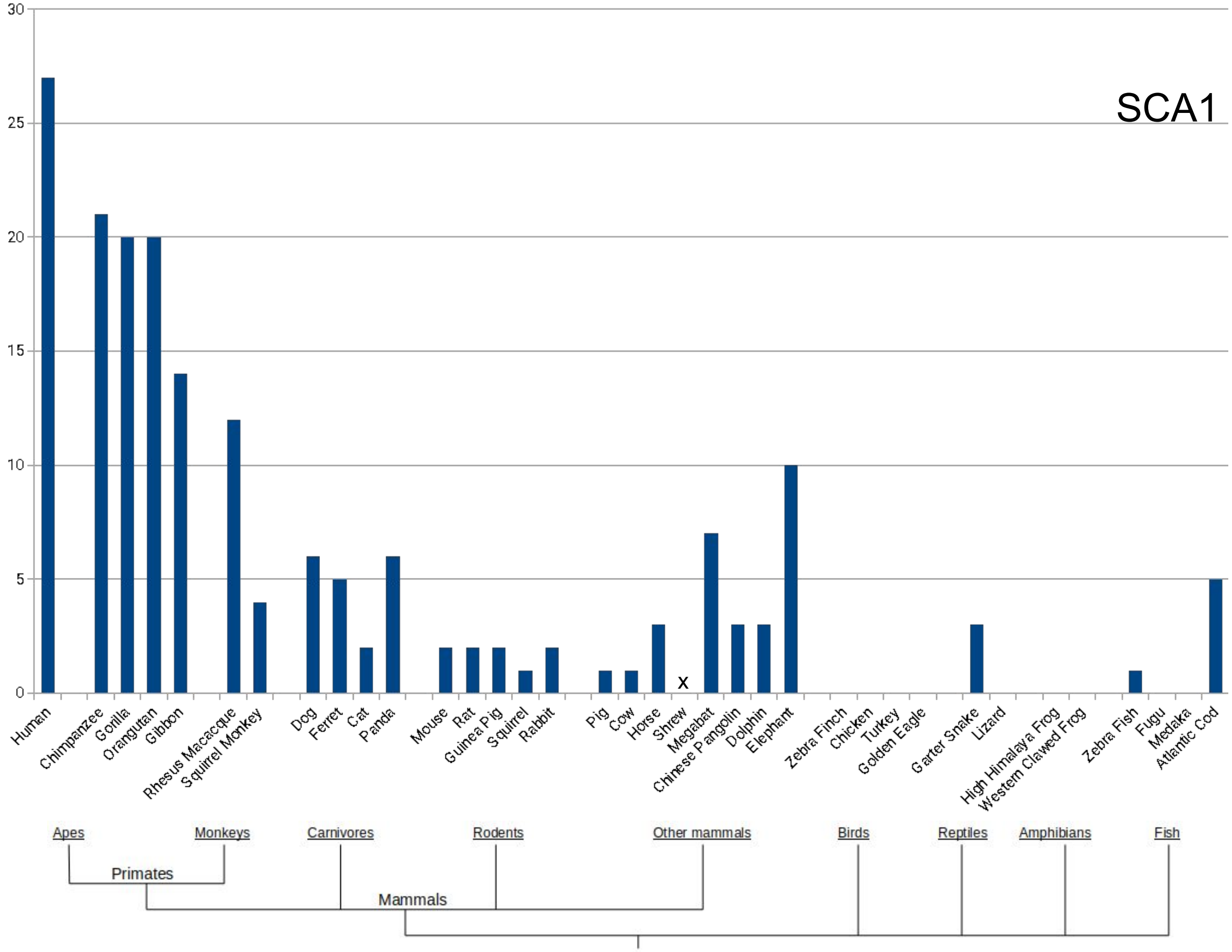
SCA1
30
25
20
15
10
5
0
x
Human
Chimpanzee
Gorilla
Orangutan
Gibbon
Rhesus Macacque
Squirrel Monkey
Dog
Ferret
Cat
Panda
Mouse
Rat
Guinea Pig
Squirrel
Rabbit
Pig
Cow
Horse
Shrew
Megabat
Chinese Pangolin
Dolphin
Elephant
Zebra Finch
Chicken
Turkey
Golden Eagle
Garter Snake
Lizard
High Himalaya Frog
Western Clawed Frog
Zebra Fish
Fugu
Medaka
Atlantic Cod
Apes
Monkeys
Carnivores
Rodents
Other mammals
Birds
Reptiles
Amphibians
Fish
Primates
Mammals

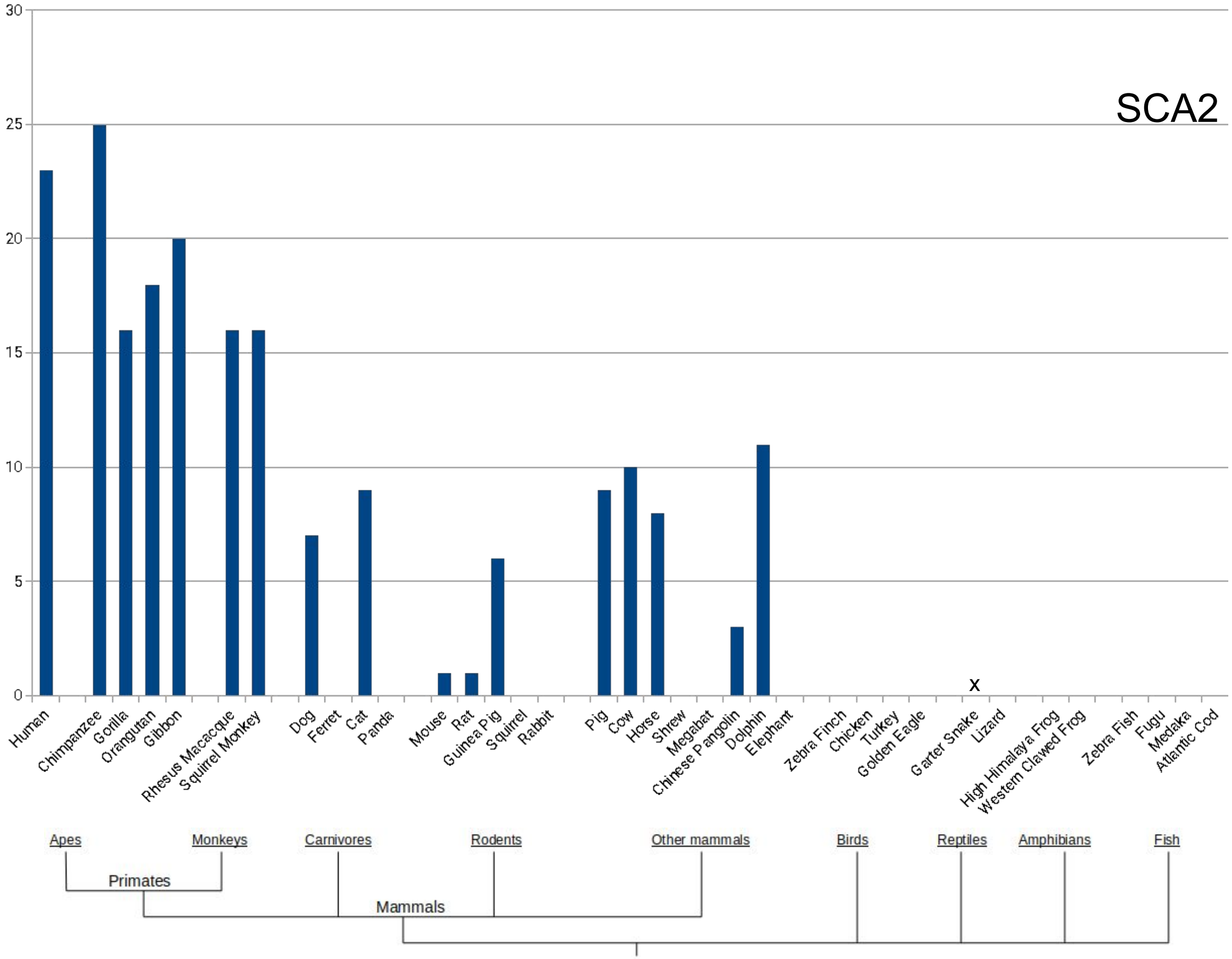
SCA2
30
25
20
15
10
5
0
Human
Chimpanzee
Gorilla
Orangutan
Gibbon
Rhesus Macacque
Squirrel Monkey
Dog
Ferret
Cat
Panda
Mouse
Rat
Guinea Pig
Squirrel
Rabbit
Pig
Cow
Horse
Shrew
Megabat
Chinese Pangolin
Dolphin
Elephant
Zebra Finch
Chicken
Turkey
Golden Eagle
Garter Snake
Lizard
High Himalaya Frog
Western Clawed Frog
Zebra Fish
Fugu
Medaka
Atlantic Cod
x
Apes
Monkeys
Carnivores
Rodents
Other mammals
Birds
Reptiles
Amphibians
Fish
Primates
Mammals

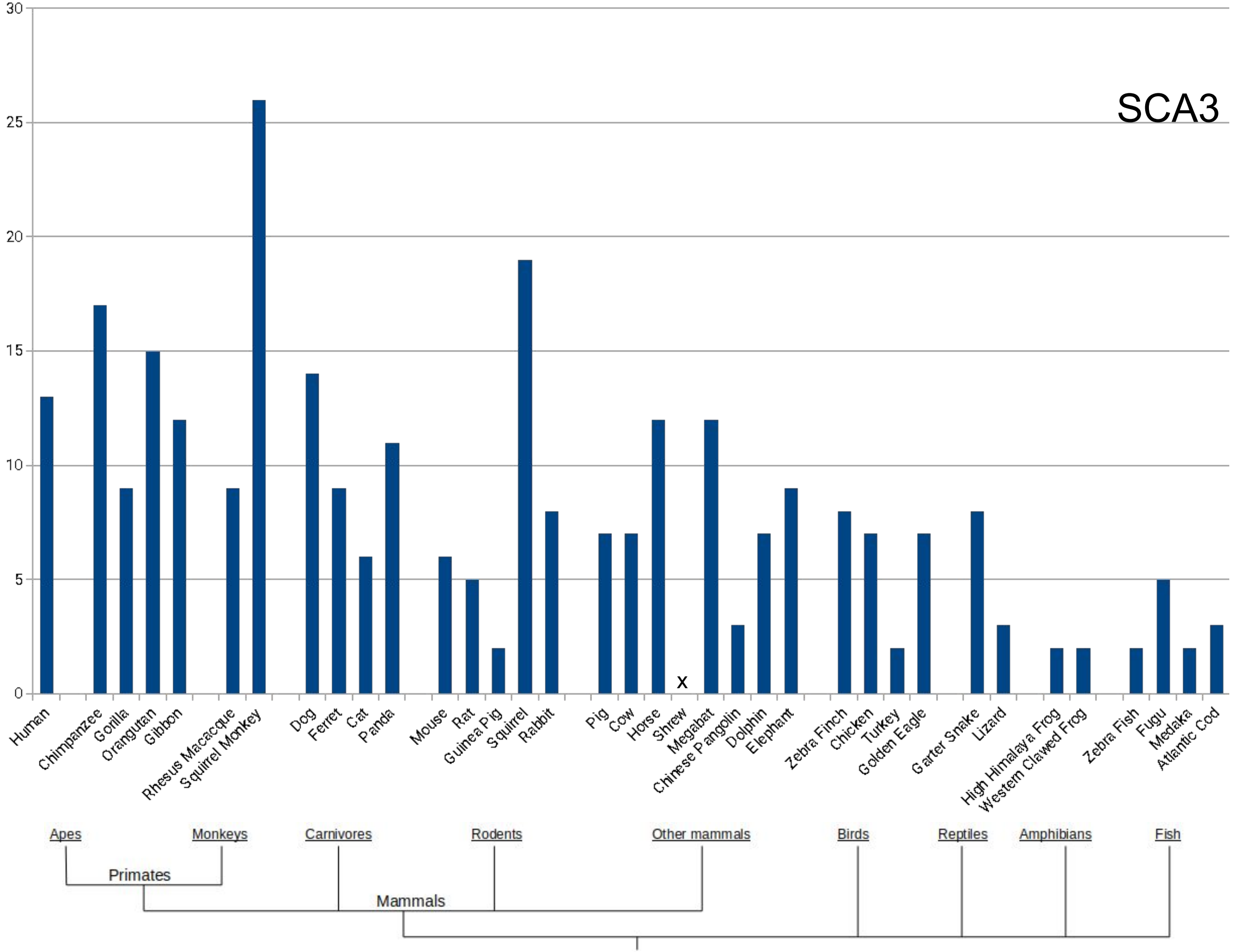
SCA3
30
25
20
15
10
5
0
x
Human
Chimpanzee
Gorilla
Orangutan
Gibbon
Rhesus Macacque
Squirrel Monkey
Dog
Ferret
Cat
Panda
Mouse
Rat
Guinea Pig
Squirrel
Rabbit
Pig
Cow
Horse
Shrew
Megabat
Chinese Pangolin
Dolphin
Elephant
Zebra Finch
Chicken
Turkey
Golden Eagle
Garter Snake
Lizard
High Himalaya Frog
Western Clawed Frog
Zebra Fish
Fugu
Medaka
Atlantic Cod
Apes
Monkeys
Carnivores
Rodents
Other mammals
Birds
Reptiles
Amphibians
Fish
Primates
Mammals

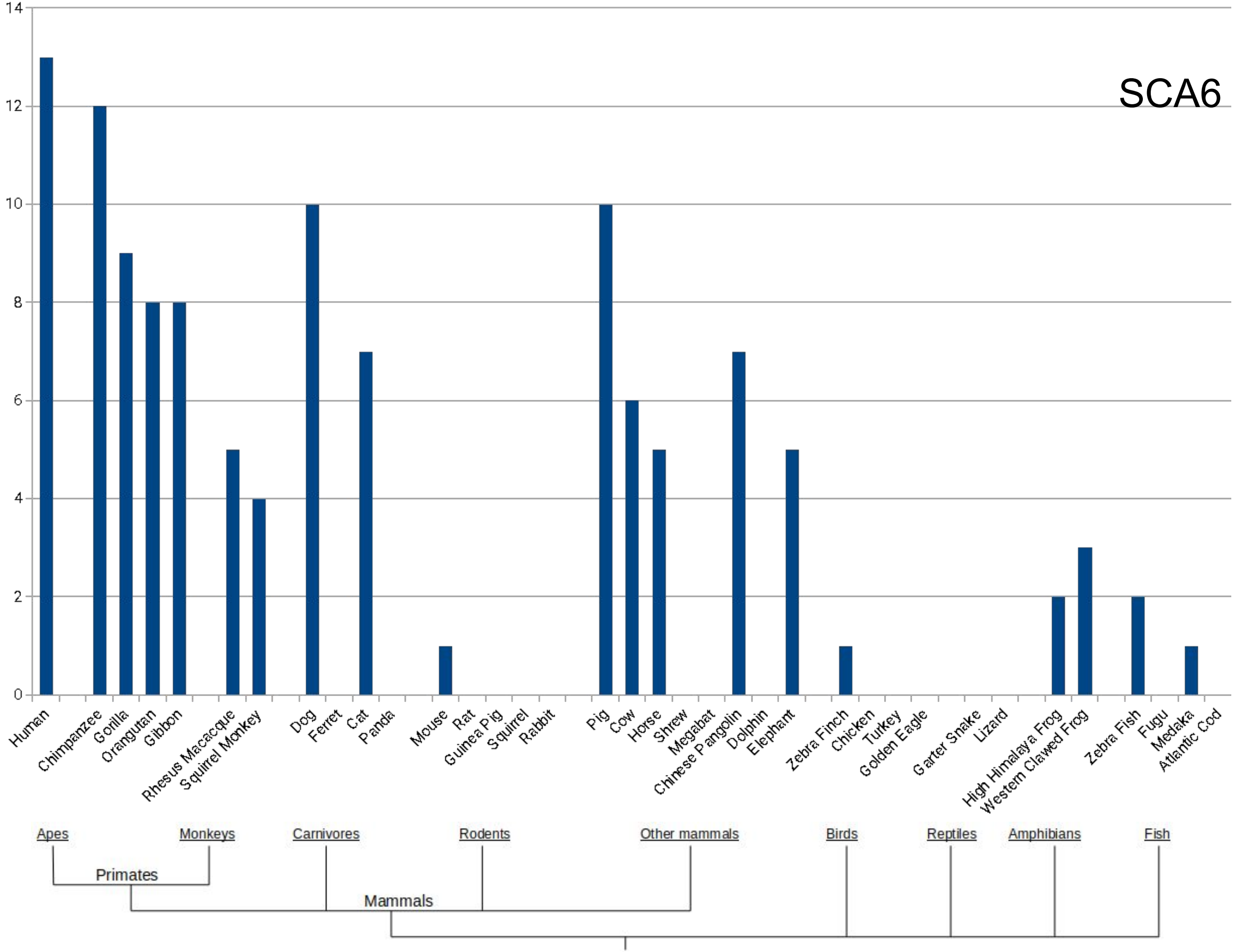

SCA6
14
12
10
8
6
4
2
0
Human
Chimpanzee
Gorilla
Orangutan
Gibbon
Rhesus Macacque
Squirrel Monkey
Dog
Ferret
Cat
Panda
Mouse
Rat
Guinea Pig
Squirrel
Rabbit
Pig
Cow
Horse
Shrew
Megabat
Chinese Pangolin
Dolphin
Elephant
Zebra Finch
Chicken
Turkey
Golden Eagle
Garter Snake
Lizard
High Himalaya Frog
Western Clawed Frog
Zebra Fish
Fugu
Medaka
Atlantic Cod
Apes
Monkeys
Carnivores
Rodents
Other mammals
Birds
Reptiles
Amphibians
Fish
Primates
Mammals

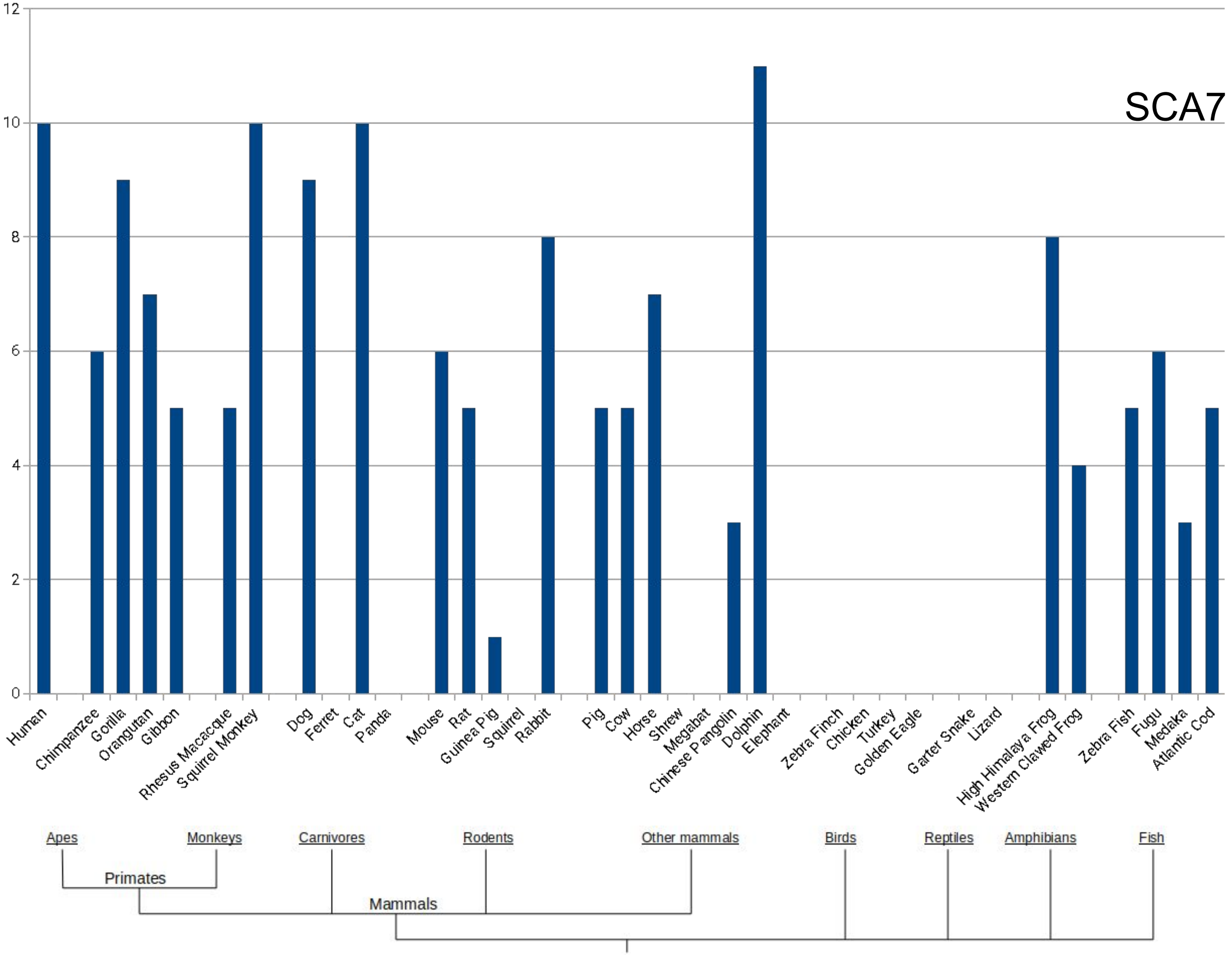

SCA7
12
10
8
6
4
2
0
Human
Chimpanzee
Gorilla
Orangutan
Gibbon
Rhesus Macacque
Squirrel Monkey
Dog
Ferret
Cat
Panda
Mouse
Rat
Guinea Pig
Squirrel
Rabbit
Pig
Cow
Horse
Shrew
Megabat
Chinese Pangolin
Dolphin
Elephant
Zebra Finch
Chicken
Turkey
Golden Eagle
Garter Snake
Lizard
High Himalaya Frog
Western Clawed Frog
Zebra Fish
Fugu
Medaka
Atlantic Cod
Apes
Monkeys
Carnivores
Rodents
Other mammals
Birds
Reptiles
Amphibians
Fish
Primates
Mammals

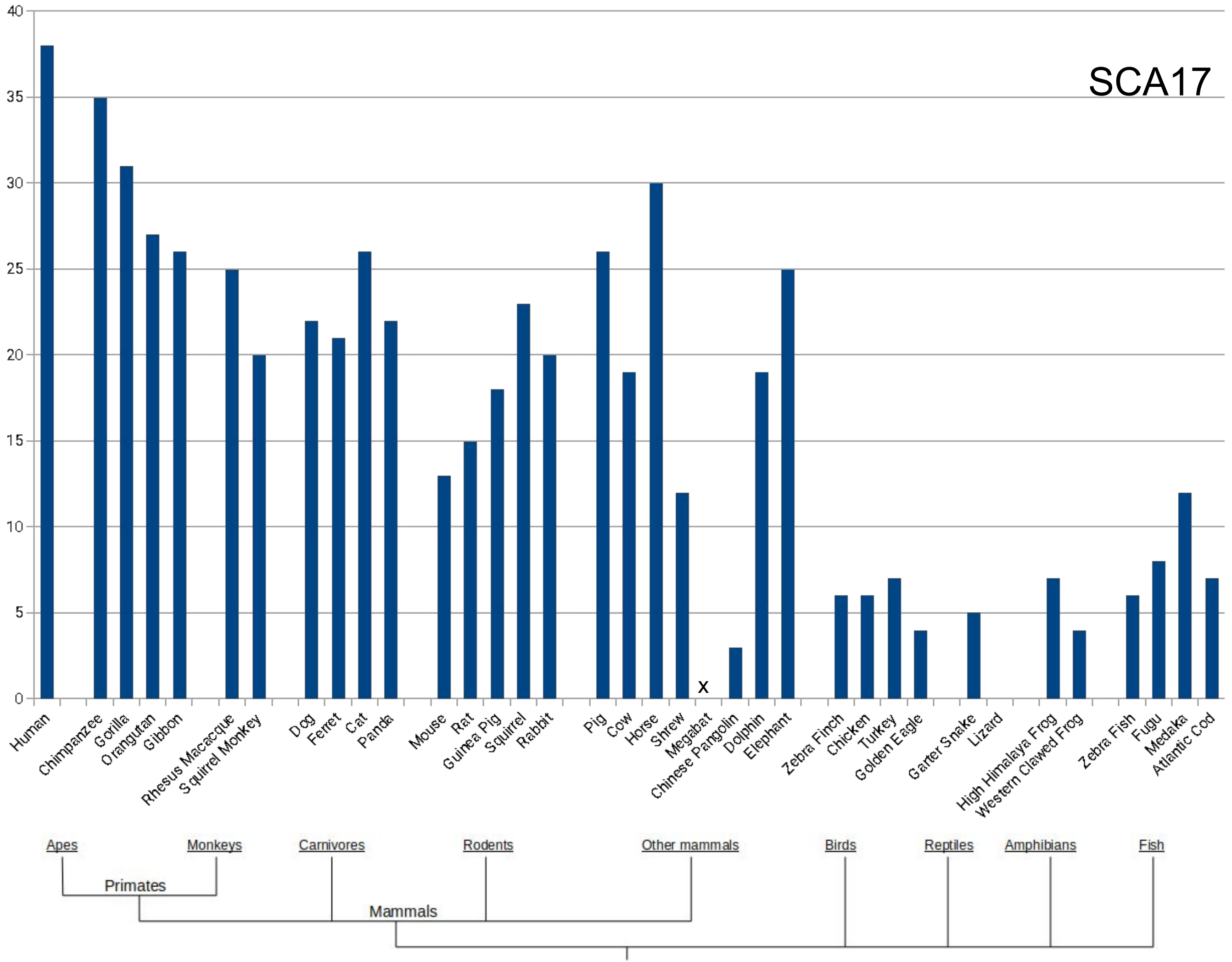
SCA17
40
35
30
25
20
15
10
5
0
x
Human
Chimpanzee
Gorilla
Orangutan
Gibbon
Rhesus Macacque
Squirrel Monkey
Dog
Ferret
Cat
Panda
Mouse
Rat
Guinea Pig
Squirrel
Rabbit
Pig
Cow
Horse
Shrew
Megabat
Chinese Pangolin
Dolphin
Elephant
Zebra Finch
Chicken
Turkey
Golden Eagle
Garter Snake
Lizard
High Himalaya Frog
Western Clawed Frog
Zebra Fish
Fugu
Medaka
Atlantic Cod
Apes
Monkeys
Carnivores
Rodents
Other mammals
Birds
Reptiles
Amphibians
Fish
Primates
Mammals

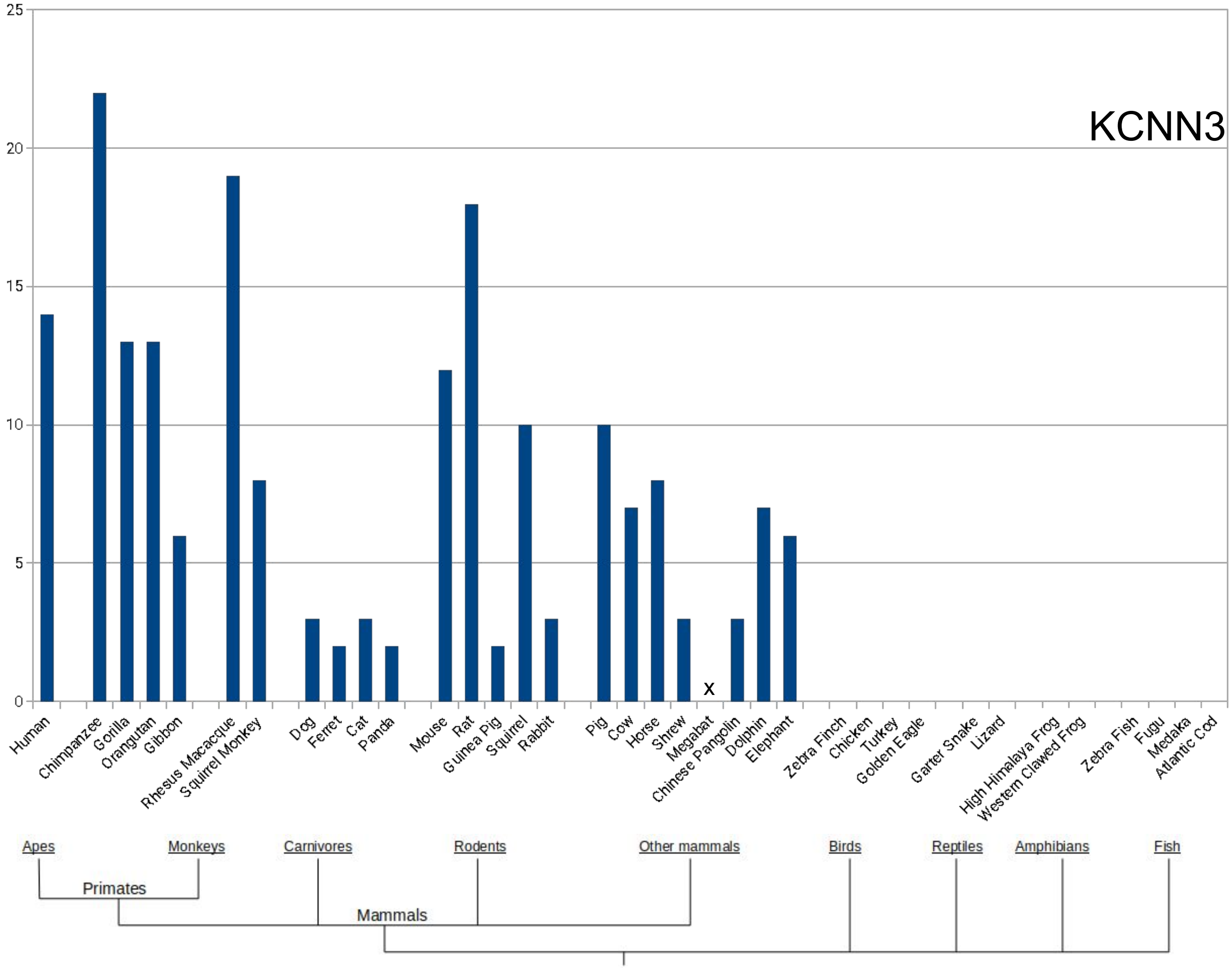
KCNN3
0
5
10
15
20
25
Human
Chimpanzee
Gorilla
Orangutan
Gibbon
Rhesus Macacque
Squirrel Monkey
Dog
Ferret
Cat
Panda
Mouse
Rat
Guinea Pig
Squirrel
Rabbit
Pig
Cow
Horse
Shrew
Megabat
Chinese Pangolin
Dolphin
Elephant
Zebra Finch
Chicken
Turkey
Golden Eagle
Garter Snake
Lizard
High Himalaya Frog
Western Clawed Frog
Zebra Fish
Fugu
Medaka
Atlantic Cod
x
Apes
Monkeys
Carnivores
Rodents
Other mammals
Birds
Reptiles
Amphibians
Fish
Primates
Mammals

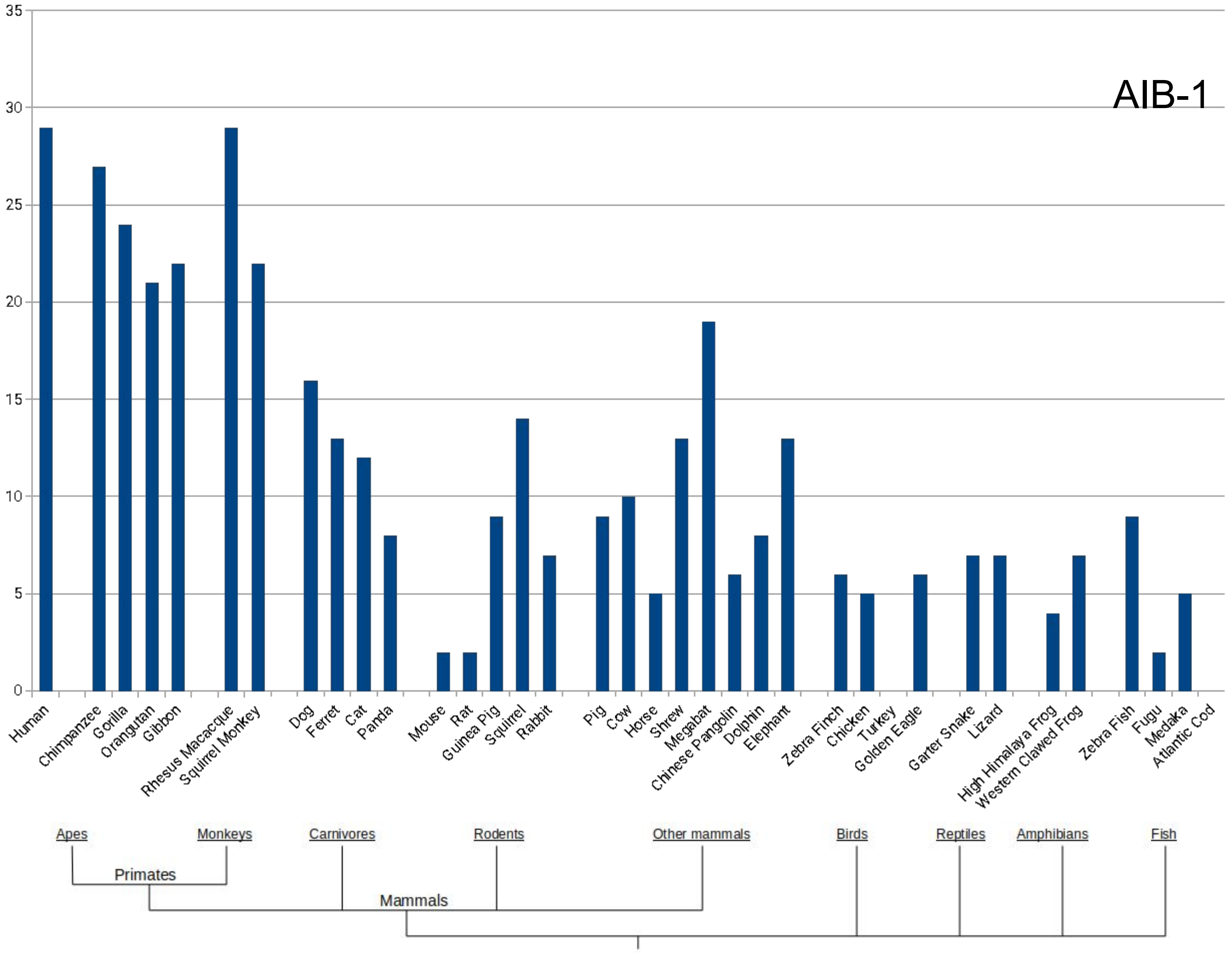
AIB-1
35
30
25
20
15
10
5
0
Human
Chimpanzee
Gorilla
Orangutan
Gibbon
Rhesus Macacque
Squirrel Monkey
Dog
Ferret
Cat
Panda
Mouse
Rat
Guinea Pig
Squirrel
Rabbit
Pig
Cow
Horse
Shrew
Megabat
Chinese Pangolin
Dolphin
Elephant
Zebra Finch
Chicken
Turkey
Golden Eagle
Garter Snake
Lizard
High Himalaya Frog
Western Clawed Frog
Zebra Fish
Fugu
Medaka
Atlantic Cod
Apes
Monkeys
Carnivores
Rodents
Other mammals
Birds
Reptiles
Amphibians
Fish
Primates
Mammals

### POLYALANINE TNR RELATED DISEASES

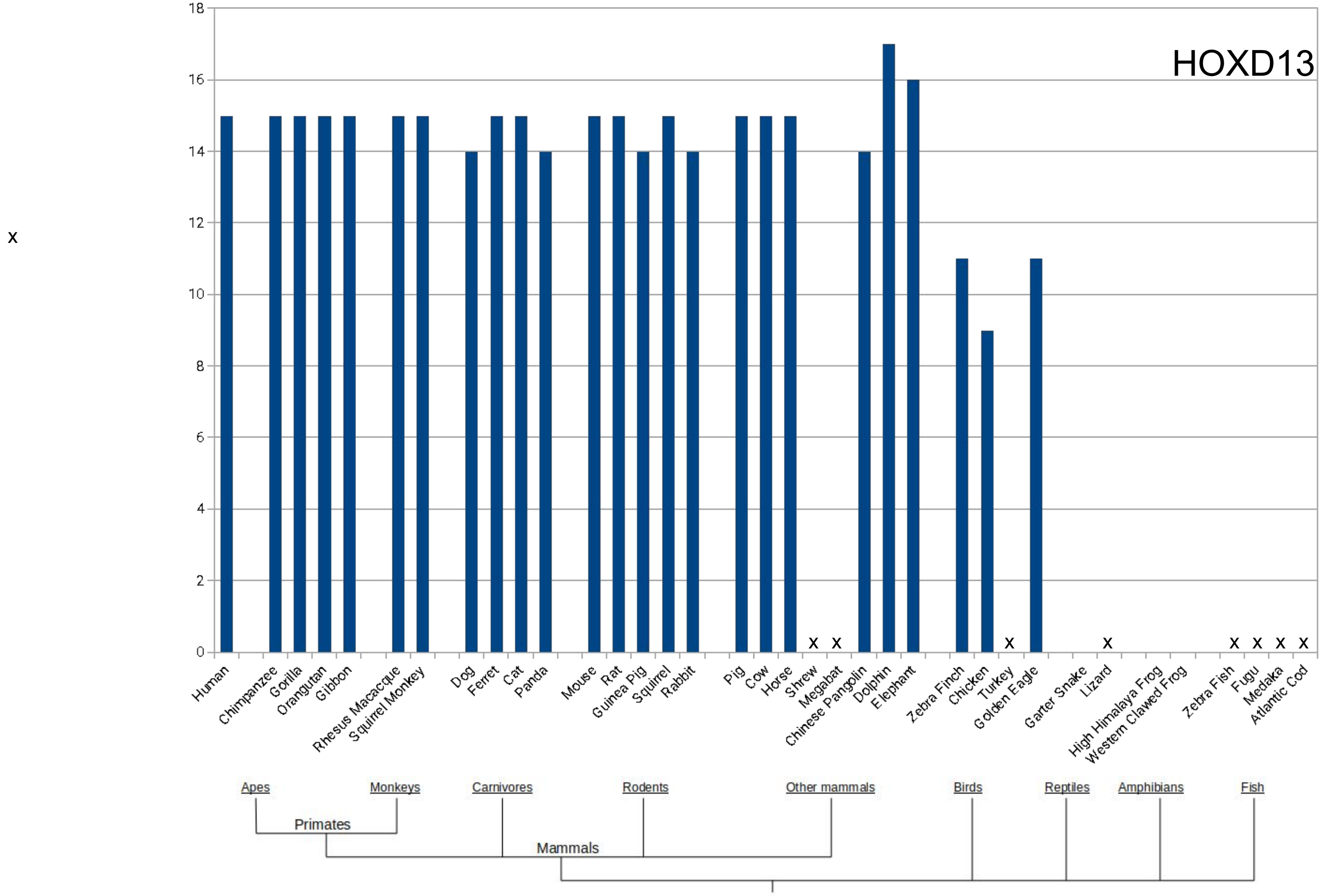
HOXD13
x
18
16
14
12
10
8
6
4
2
0
x
x
x
x
x
x
x
x
Human
Chimpanzee
Gorilla
Orangutan
Gibbon
Rhesus Macacque
Squirrel Monkey
Dog
Ferret
Cat
Panda
Mouse
Rat
Guinea Pig
Squirrel
Rabbit
Pig
Cow
Horse
Shrew
Megabat
Chinese Pangolin
Dolphin
Elephant
Zebra Finch
Chicken
Turkey
Golden Eagle
Garter Snake
Lizard
High Himalaya Frog
Western Clawed Frog
Zebra Fish
Fugu
Medaka
Atlantic Cod
Apes
Monkeys
Carnivores
Rodents
Other mammals
Birds
Reptiles
Amphibians
Fish
Primates
Mammals

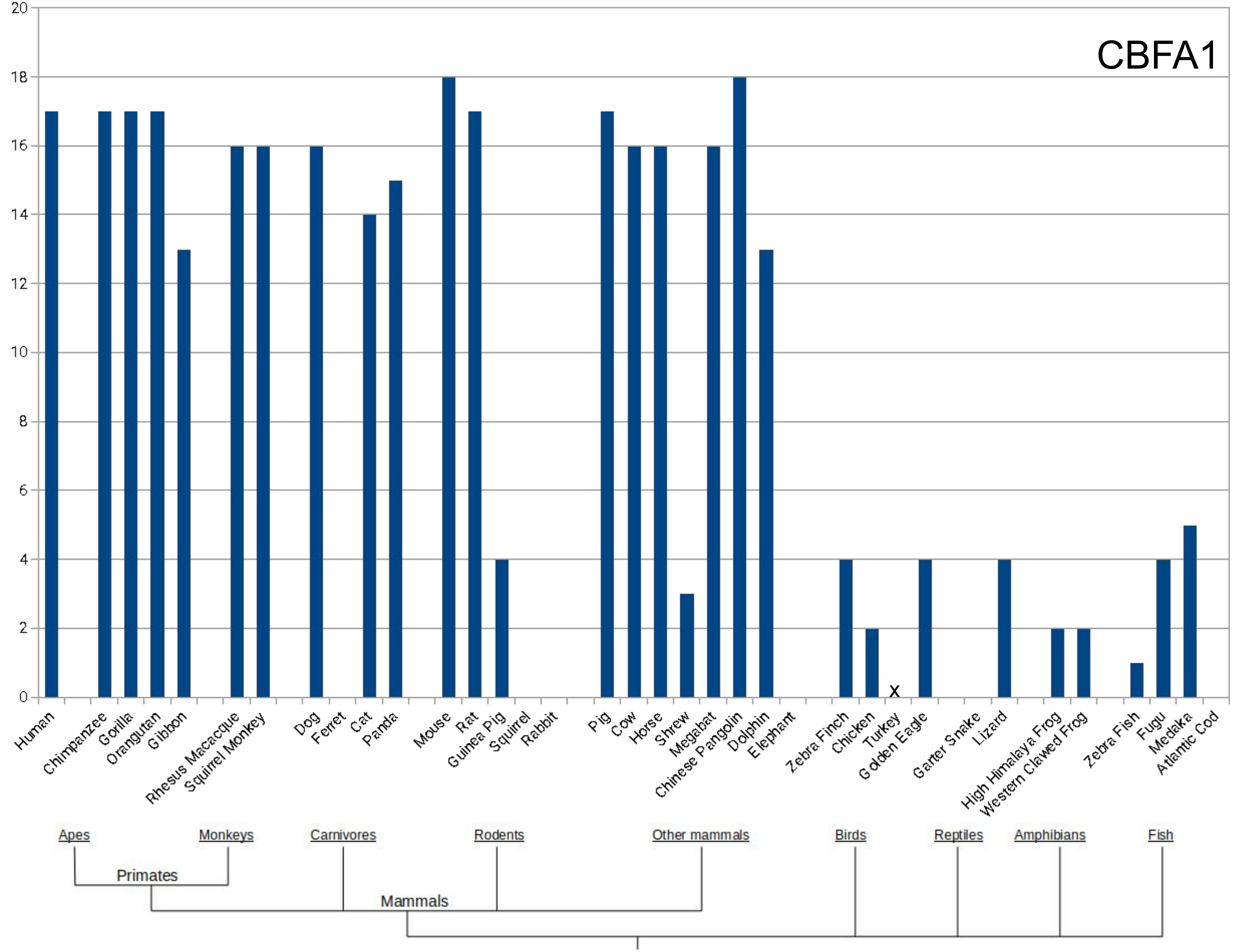
CBFA1
0
2
4
6
8
10
12
14
16
18
20
Human
Chimpanzee
Gorilla
Orangutan
Gibbon
Rhesus Macacque
Squirrel Monkey
Dog
Ferret
Cat
Panda
Mouse
Rat
Guinea Pig
Squirrel
Rabbit
Pig
Cow
Horse
Shrew
Megabat
Chinese Pangolin
Dolphin
Elephant
Zebra Finch
Chicken
Turkey
Golden Eagle
Garter Snake
Lizard
High Himalaya Frog
Western Clawed Frog
Zebra Fish
Fugu
Medaka
Atlantic Cod
x
Apes
Monkeys
Carnivores
Rodents
Other mammals
Birds
Reptiles
Amphibians
Fish
Primates
Mammals

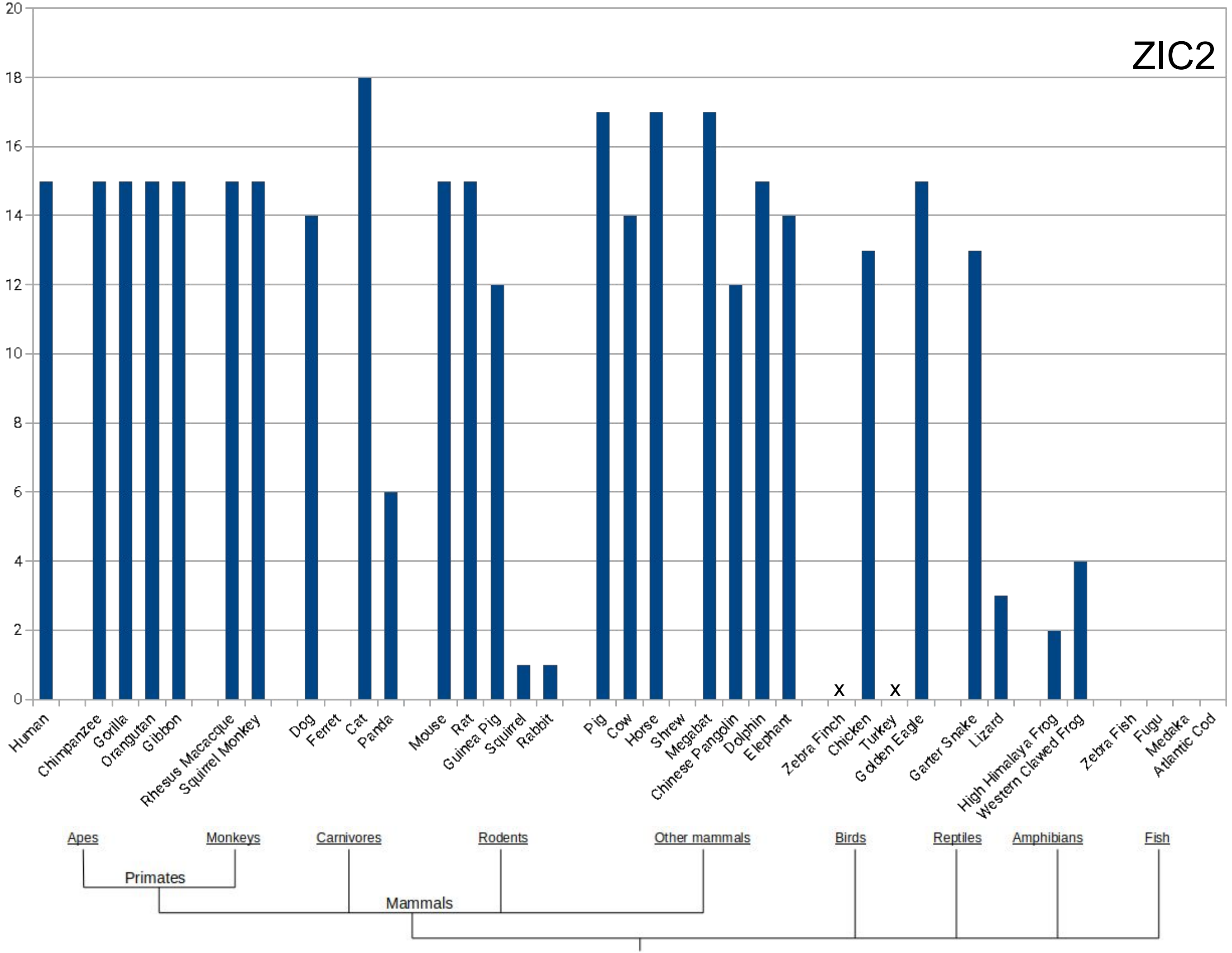
ZIC2
20
18
16
14
12
10
8
6
4
2
0
x
x
Human
Chimpanzee
Gorilla
Orangutan
Gibbon
Rhesus Macacque
Squirrel Monkey
Dog
Ferret
Cat
Panda
Mouse
Rat
Guinea Pig
Squirrel
Rabbit
Pig
Cow
Horse
Shrew
Megabat
Chinese Pangolin
Dolphin
Elephant
Zebra Finch
Chicken
Turkey
Golden Eagle
Garter Snake
Lizard
High Himalaya Frog
Western Clawed Frog
Zebra Fish
Fugu
Medaka
Atlantic Cod
Apes
Monkeys
Carnivores
Rodents
Other mammals
Birds
Reptiles
Amphibians
Fish
Primates
Mammals

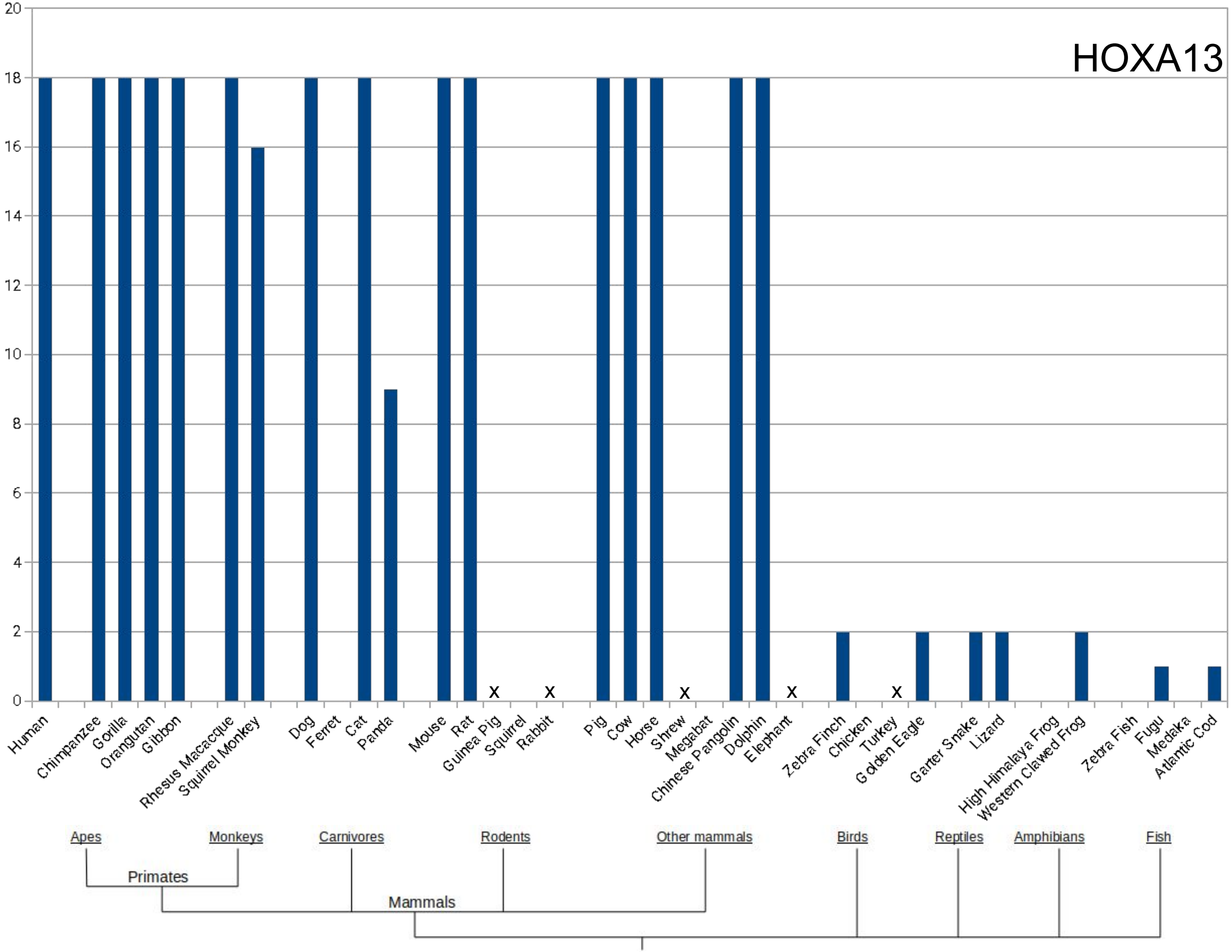
HOXA13
20
18
16
14
12
10
8
6
4
2
0
x
x
x
x
x
Human
Chimpanzee
Gorilla
Orangutan
Gibbon
Rhesus Macacque
Squirrel Monkey
Dog
Ferret
Cat
Panda
Mouse
Rat
Guinea Pig
Squirrel
Rabbit
Pig
Cow
Horse
Shrew
Megabat
Chinese Pangolin
Dolphin
Elephant
Zebra Finch
Chicken
Turkey
Golden Eagle
Garter Snake
Lizard
High Himalaya Frog
Western Clawed Frog
Zebra Fish
Fugu
Medaka
Atlantic Cod
Apes
Monkeys
Carnivores
Rodents
Other mammals
Birds
Reptiles
Amphibians
Fish
Primates
Mammals

x

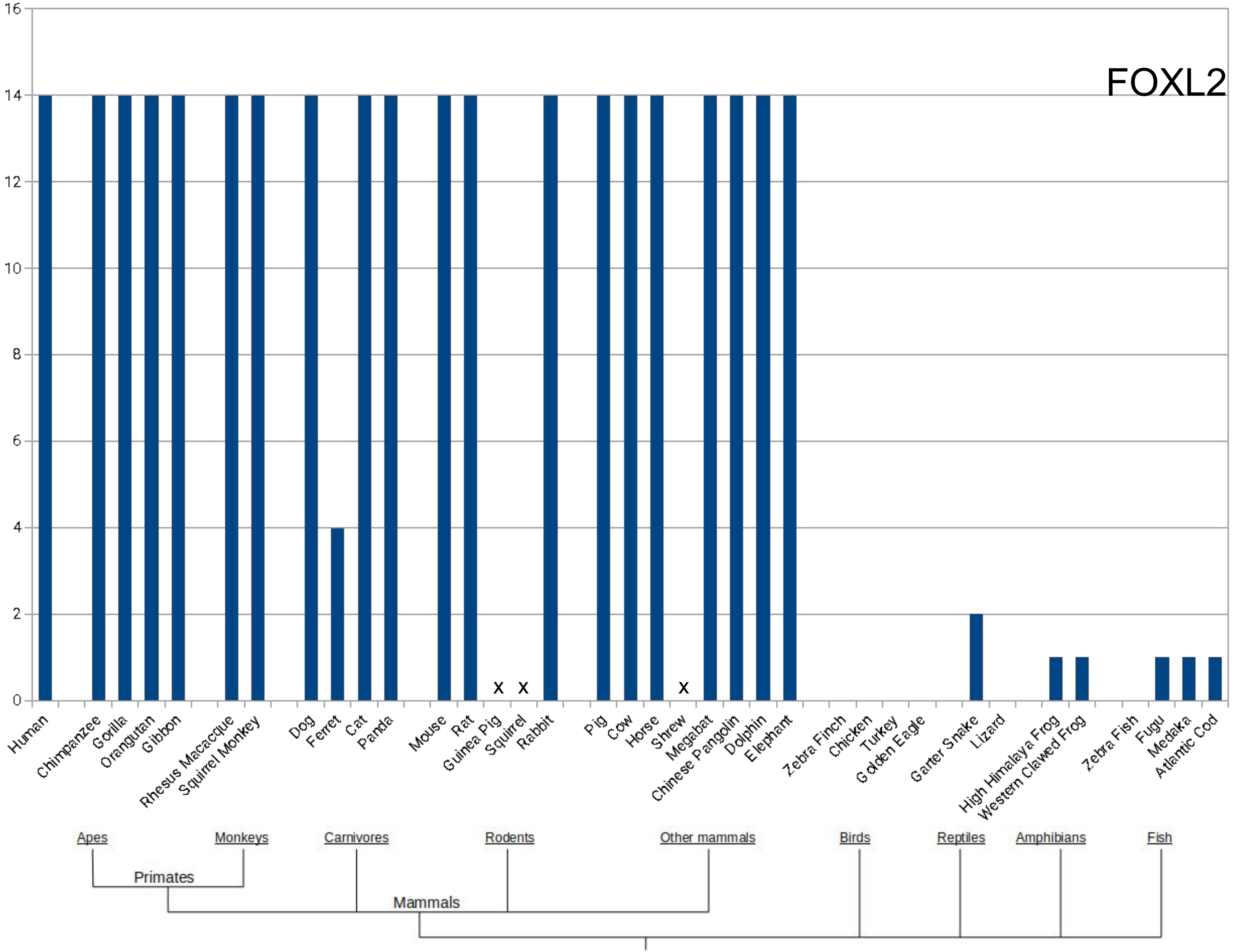

FOXL2
16
14
12
10
8
6
4
2
0
x
x
x
Human
Chimpanzee
Gorilla
Orangutan
Gibbon
Rhesus Macacque
Squirrel Monkey
Dog
Ferret
Cat
Panda
Mouse
Rat
Guinea Pig
Squirrel
Rabbit
Pig
Cow
Horse
Shrew
Megabat
Chinese Pangolin
Dolphin
Elephant
Zebra Finch
Chicken
Turkey
Golden Eagle
Garter Snake
Lizard
High Himalaya Frog
Western Clawed Frog
Zebra Fish
Fugu
Medaka
Atlantic Cod
Apes
Monkeys
Carnivores
Rodents
Other mammals
Birds
Reptiles
Amphibians
Fish
Primates
Mammals

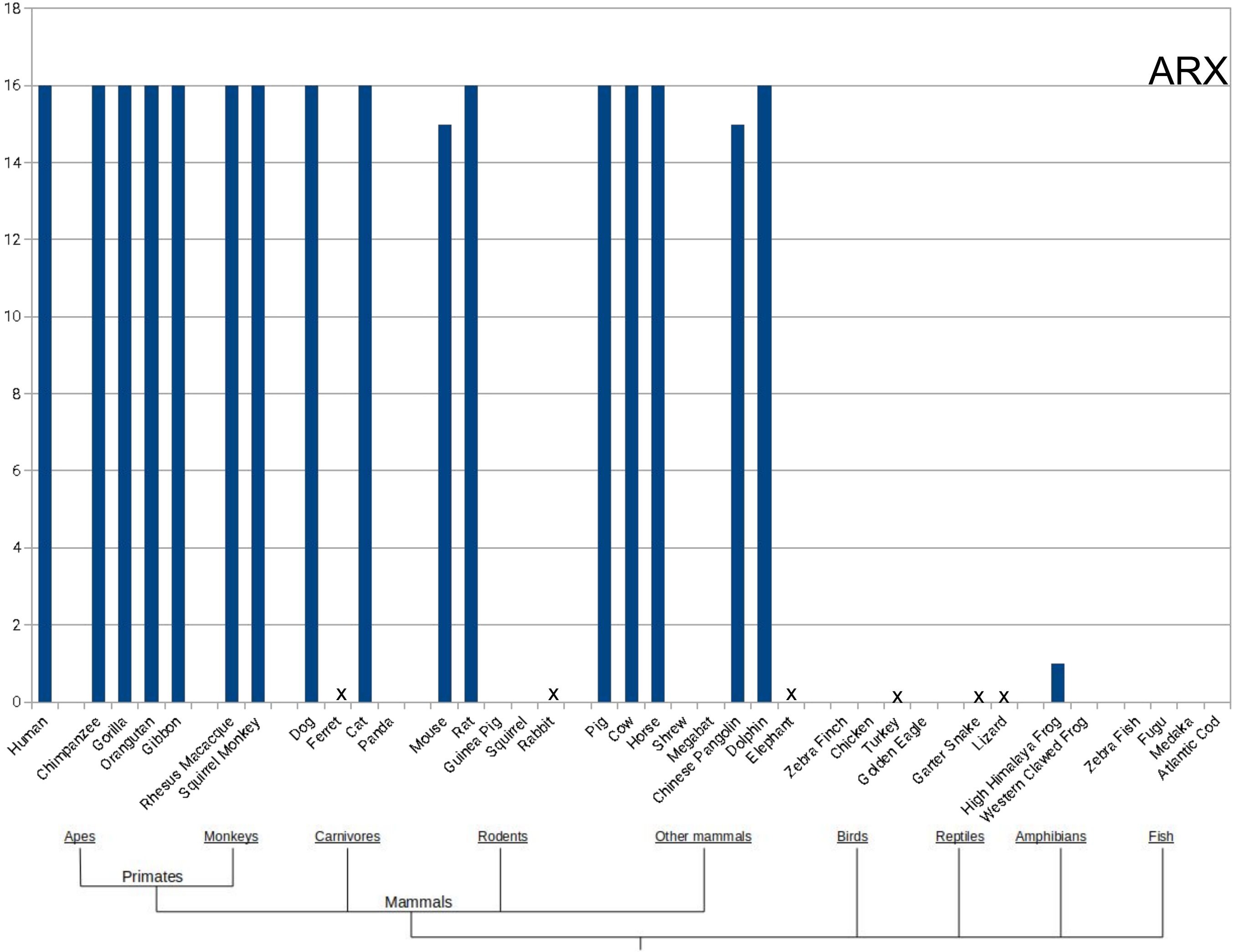
ARX
18
16
14
12
10
8
6
4
2
0
x
x
x
x
x
x
Human
Chimpanzee
Gorilla
Orangutan
Gibbon
Rhesus Macacque
Squirrel Monkey
Dog
Ferret
Cat
Panda
Mouse
Rat
Guinea Pig
Squirrel
Rabbit
Pig
Cow
Horse
Shrew
Megabat
Chinese Pangolin
Dolphin
Elephant
Zebra Finch
Chicken
Turkey
Golden Eagle
Garter Snake
Lizard
High Himalaya Frog
Western Clawed Frog
Zebra Fish
Fugu
Medaka
Atlantic Cod
Apes
Monkeys
Carnivores
Rodents
Other mammals
Birds
Reptiles
Amphibians
Fish
Primates
Mammals

### POLYASPARTATE TNR RELATED DISEASE

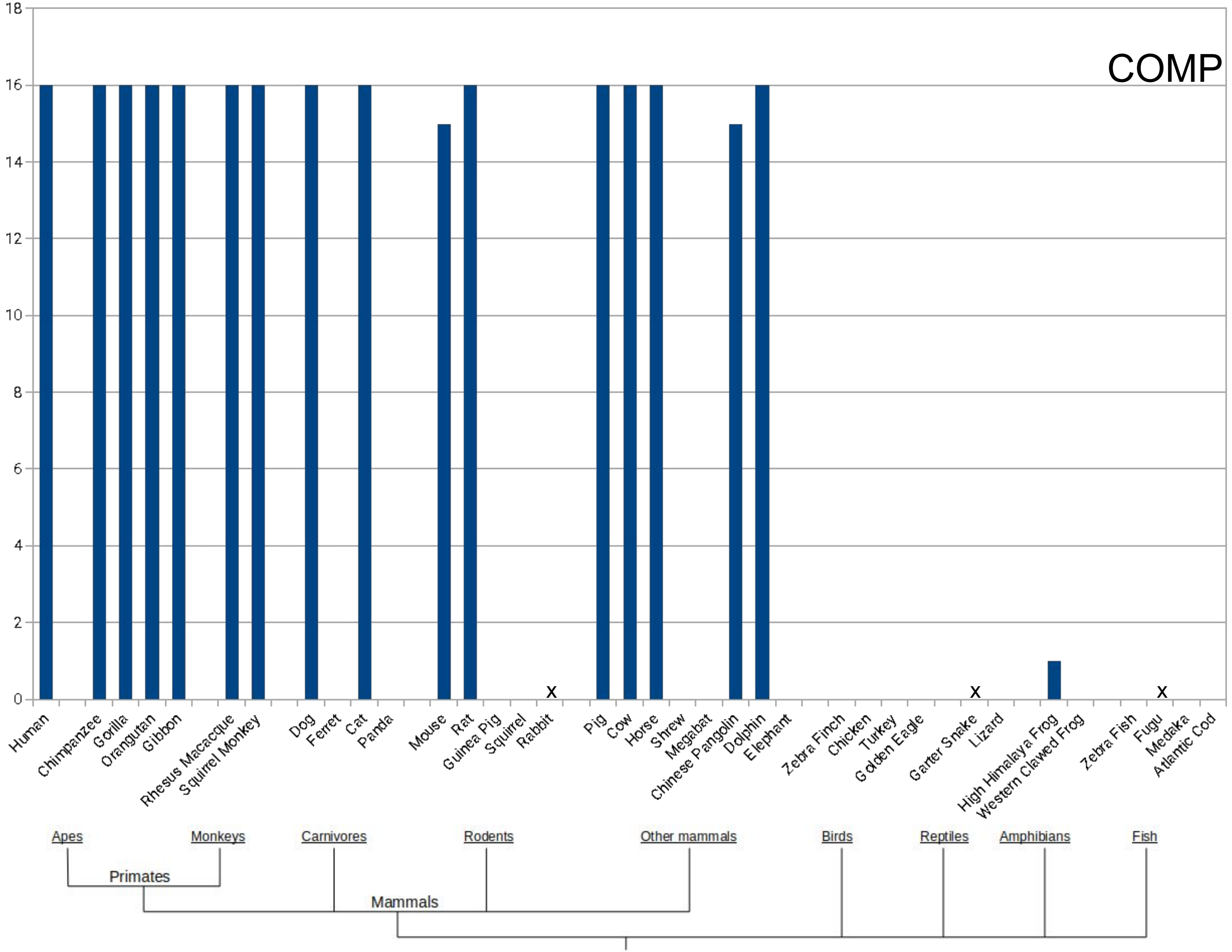
COMP
18
16
14
12
10
8
6
4
2
0
Human
Chimpanzee
Gorilla
Orangutan
Gibbon
Rhesus Macacque
Squirrel Monkey
Dog
Ferret
Cat
Panda
Mouse
Rat
Guinea Pig
Squirrel
Rabbit
Pig
Cow
Horse
Shrew
Megabat
Chinese Pangolin
Dolphin
Elephant
Zebra Finch
Chicken
Turkey
Golden Eagle
Garter Snake
Lizard
High Himalaya Frog
Western Clawed Frog
Zebra Fish
Fugu
Medaka
Atlantic Cod
x
x
x
Apes
Monkeys
Carnivores
Rodents
Other mammals
Birds
Reptiles
Amphibians
Fish
Primates
Mammals

### NON CODING REGION TNR RELATED DISEASES

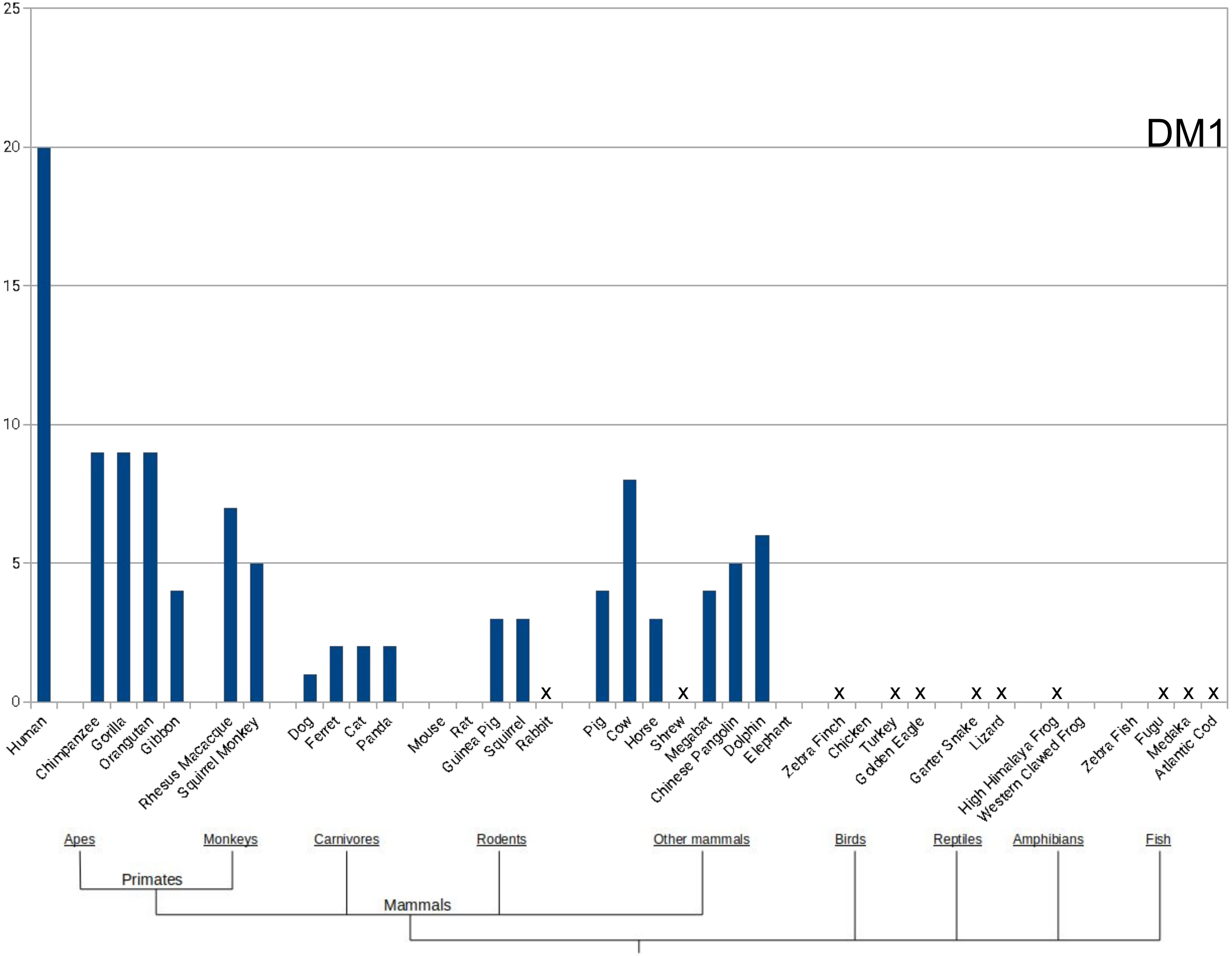
DM1
25
20
15
10
5
0
Human
Chimpanzee
Gorilla
Orangutan
Gibbon
Rhesus Macacque
Squirrel Monkey
Dog
Ferret
Cat
Panda
Mouse
Rat
Guinea Pig
Squirrel
Rabbit
Pig
Cow
Horse
Shrew
Megabat
Chinese Pangolin
Dolphin
Elephant
Zebra Finch
Chicken
Turkey
Golden Eagle
Garter Snake
Lizard
High Himalaya Frog
Western Clawed Frog
Zebra Fish
Fugu
Medaka
Atlantic Cod
Apes
Monkeys
Carnivores
Rodents
Other mammals
Birds
Reptiles
Amphibians
Fish
Primates
Mammals

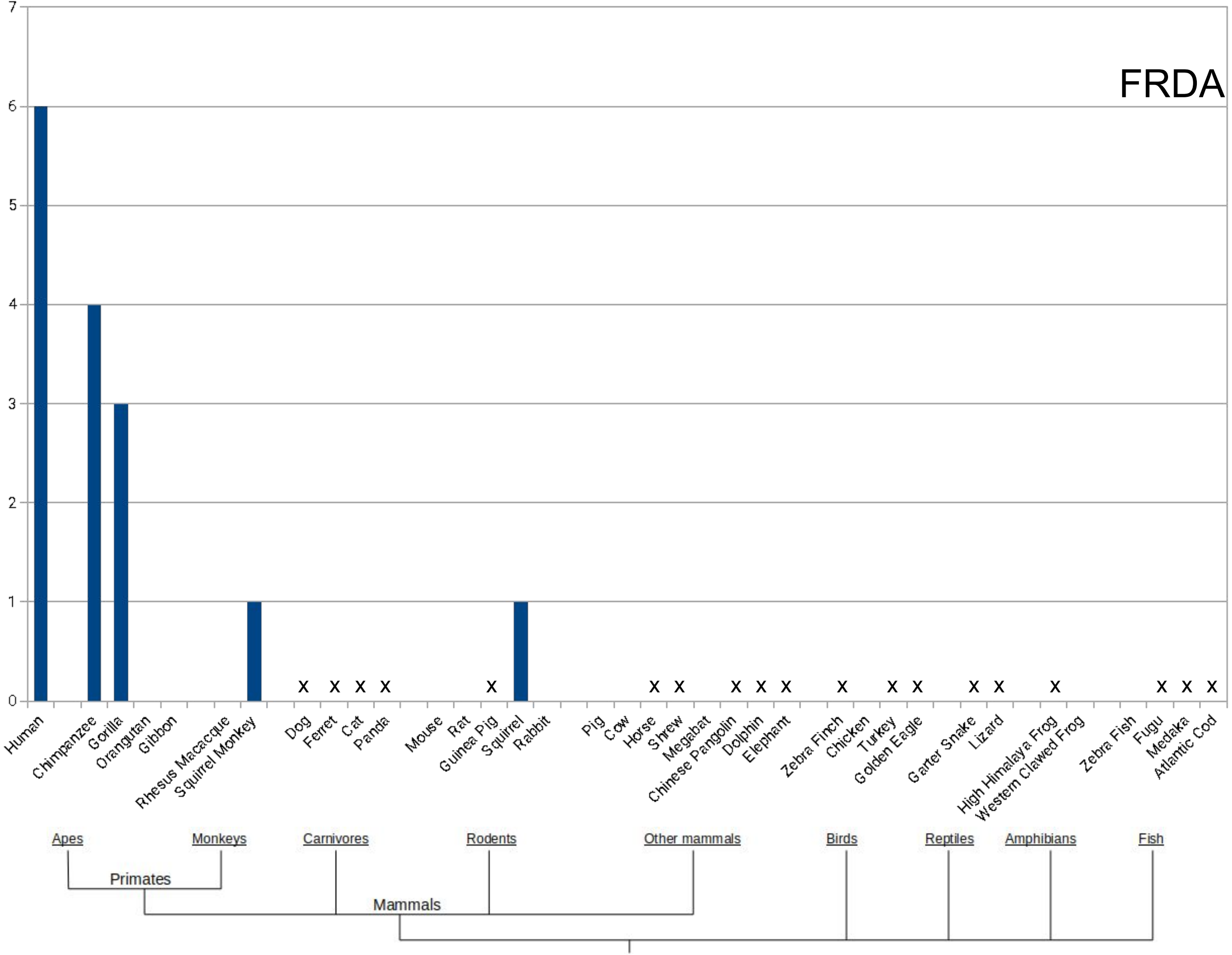
FRDA
0
1
2
3
4
5
6
7
Human
Chimpanzee
Gorilla
Orangutan
Gibbon
Rhesus Macacque
Squirrel Monkey
Dog
Ferret
Cat
Panda
Mouse
Rat
Guinea Pig
Squirrel
Rabbit
Pig
Cow
Horse
Shrew
Megabat
Chinese Pangolin
Dolphin
Elephant
Zebra Finch
Chicken
Turkey
Golden Eagle
Garter Snake
Lizard
High Himalaya Frog
Western Clawed Frog
Zebra Fish
Fugu
Medaka
Atlantic Cod
Apes
Monkeys
Carnivores
Rodents
Other mammals
Birds
Reptiles
Amphibians
Fish
Primates
Mammals

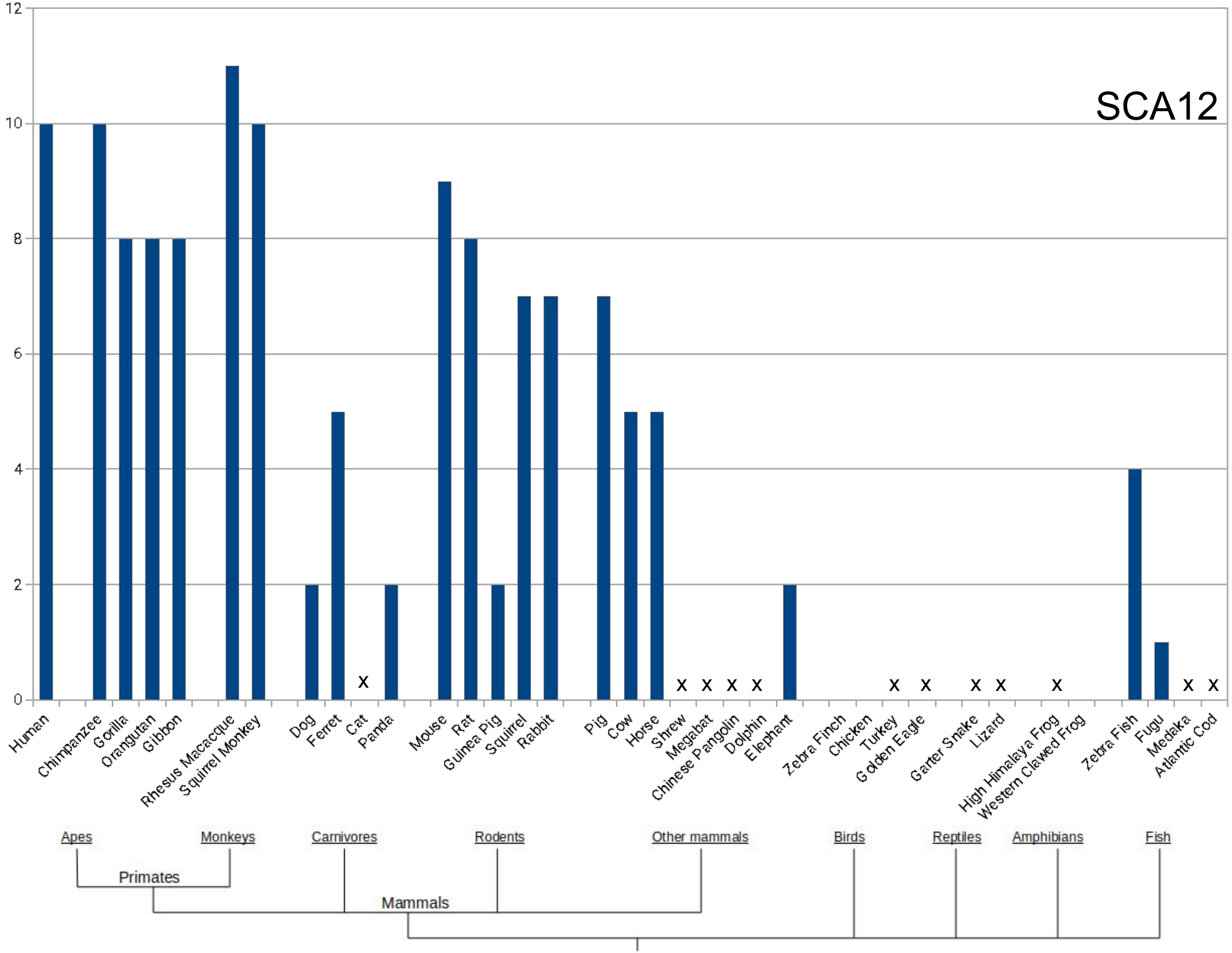
SCA12
12
10
8
6
4
2
0
Human
Chimpanzee
Gorilla
Orangutan
Gibbon
Rhesus Macacque
Squirrel Monkey
Dog
Ferret
Cat
Panda
Mouse
Rat
Guinea Pig
Squirrel
Rabbit
Pig
Cow
Horse
Shrew
Megabat
Chinese Pangolin
Dolphin
Elephant
Zebra Finch
Chicken
Turkey
Golden Eagle
Garter Snake
Lizard
High Himalaya Frog
Western Clawed Frog
Zebra Fish
Fugu
Medaka
Atlantic Cod
Apes
Monkeys
Carnivores
Rodents
Other mammals
Birds
Reptiles
Amphibians
Fish
Primates
Mammals

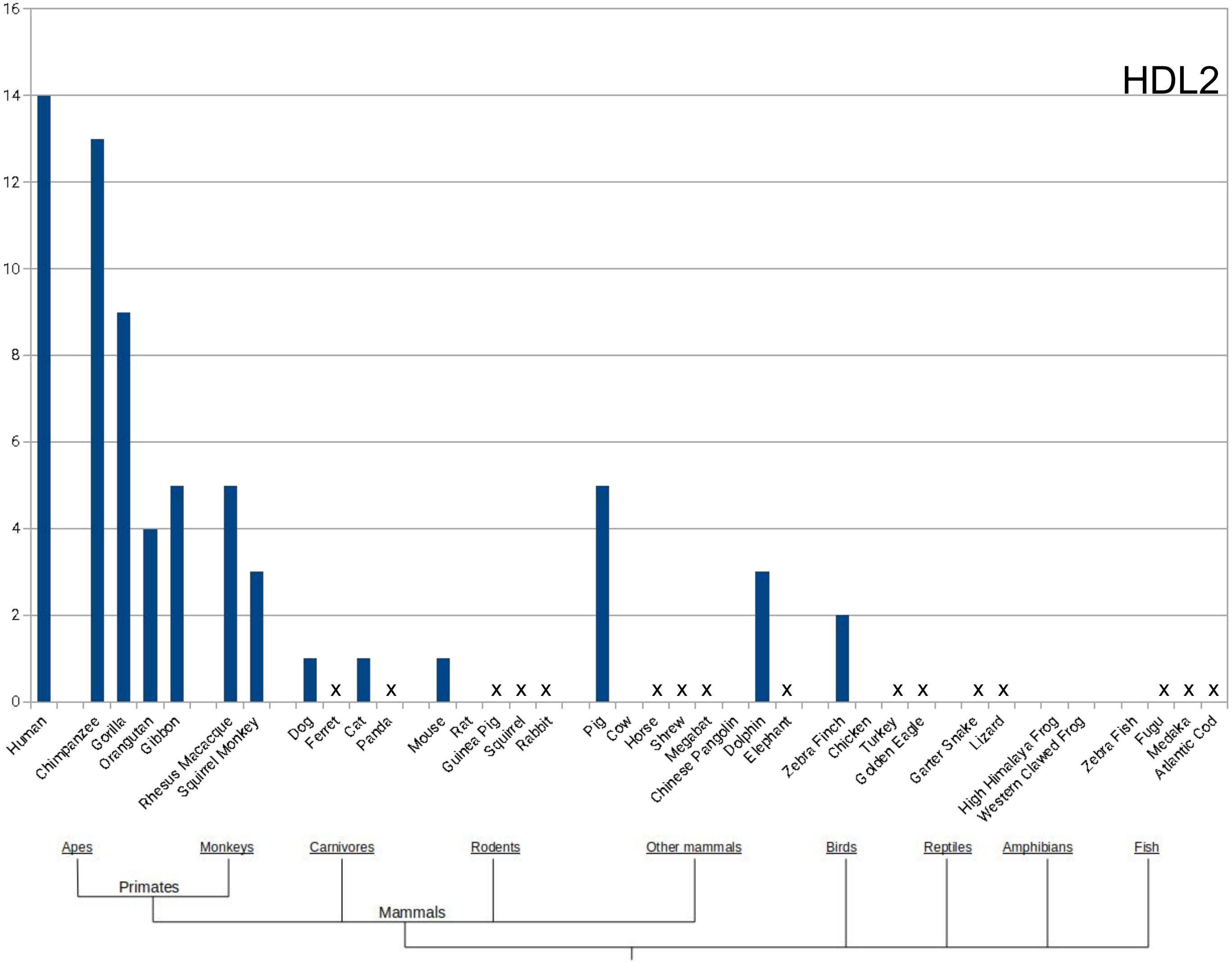
HDL2
16
14
12
10
8
6
4
2
0
Human
Chimpanzee
Gorilla
Orangutan
Gibbon
Rhesus Macacque
Squirrel Monkey
Dog
Ferret
Cat
Panda
Mouse
Rat
Guinea Pig
Squirrel
Rabbit
Pig
Cow
Horse
Shrew
Megabat
Chinese Pangolin
Dolphin
Elephant
Zebra Finch
Chicken
Turkey
Golden Eagle
Garter Snake
Lizard
High Himalaya Frog
Western Clawed Frog
Zebra Fish
Fugu
Medaka
Atlantic Cod
Apes
Monkeys
Primates
Carnivores
Rodents
Other mammals
Mammals
Birds
Reptiles
Amphibians
Fish

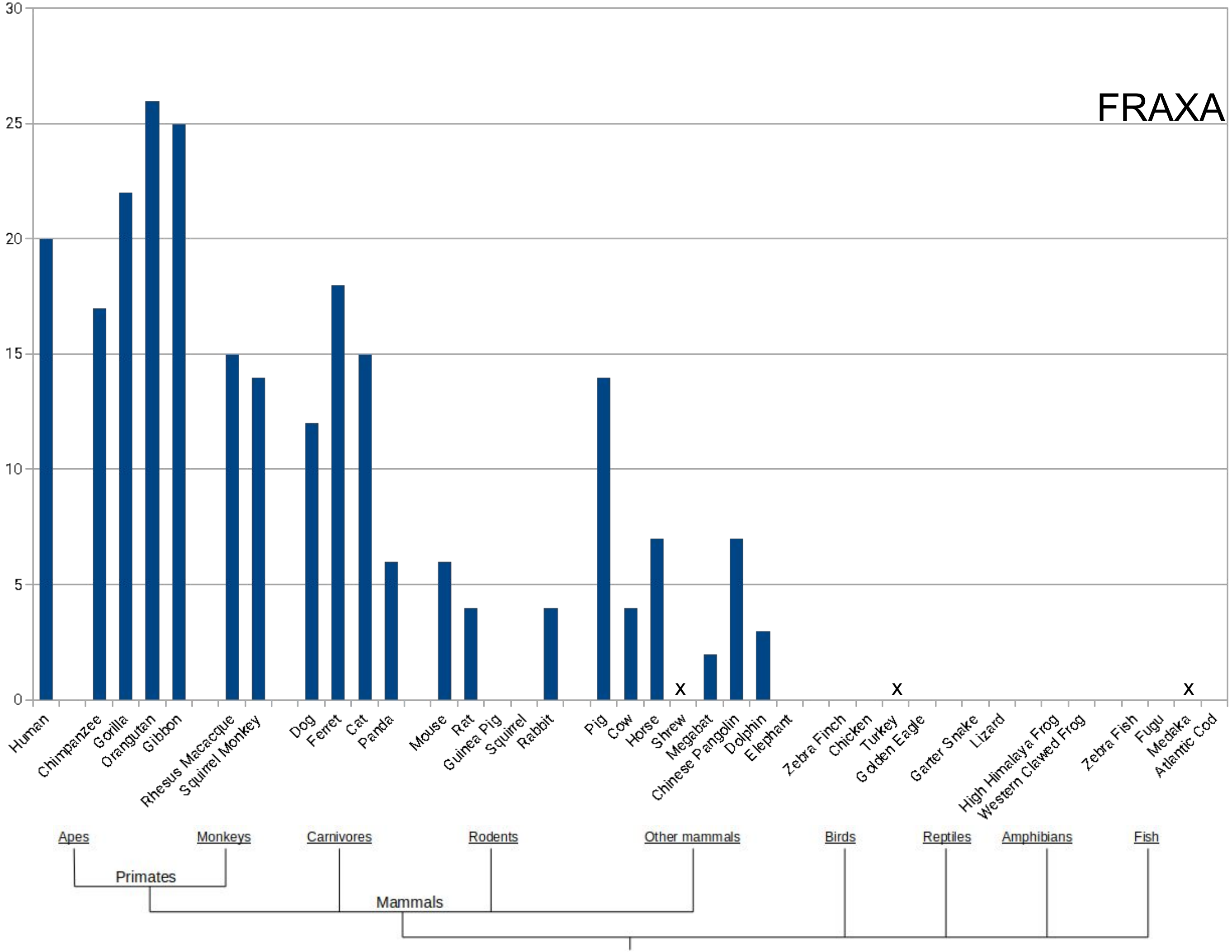
FRAXA
30
25
20
15
10
5
0
x
x
x
Human
Chimpanzee
Gorilla
Orangutan
Gibbon
Rhesus Macacque
Squirrel Monkey
Dog
Ferret
Cat
Panda
Mouse
Rat
Guinea Pig
Squirrel
Rabbit
Pig
Cow
Horse
Shrew
Megabat
Chinese Pangolin
Dolphin
Elephant
Zebra Finch
Chicken
Turkey
Golden Eagle
Garter Snake
Lizard
High Himalaya Frog
Western Clawed Frog
Zebra Fish
Fugu
Medaka
Atlantic Cod
Apes
Monkeys
Carnivores
Rodents
Other mammals
Birds
Reptiles
Amphibians
Fish
Primates
Mammals

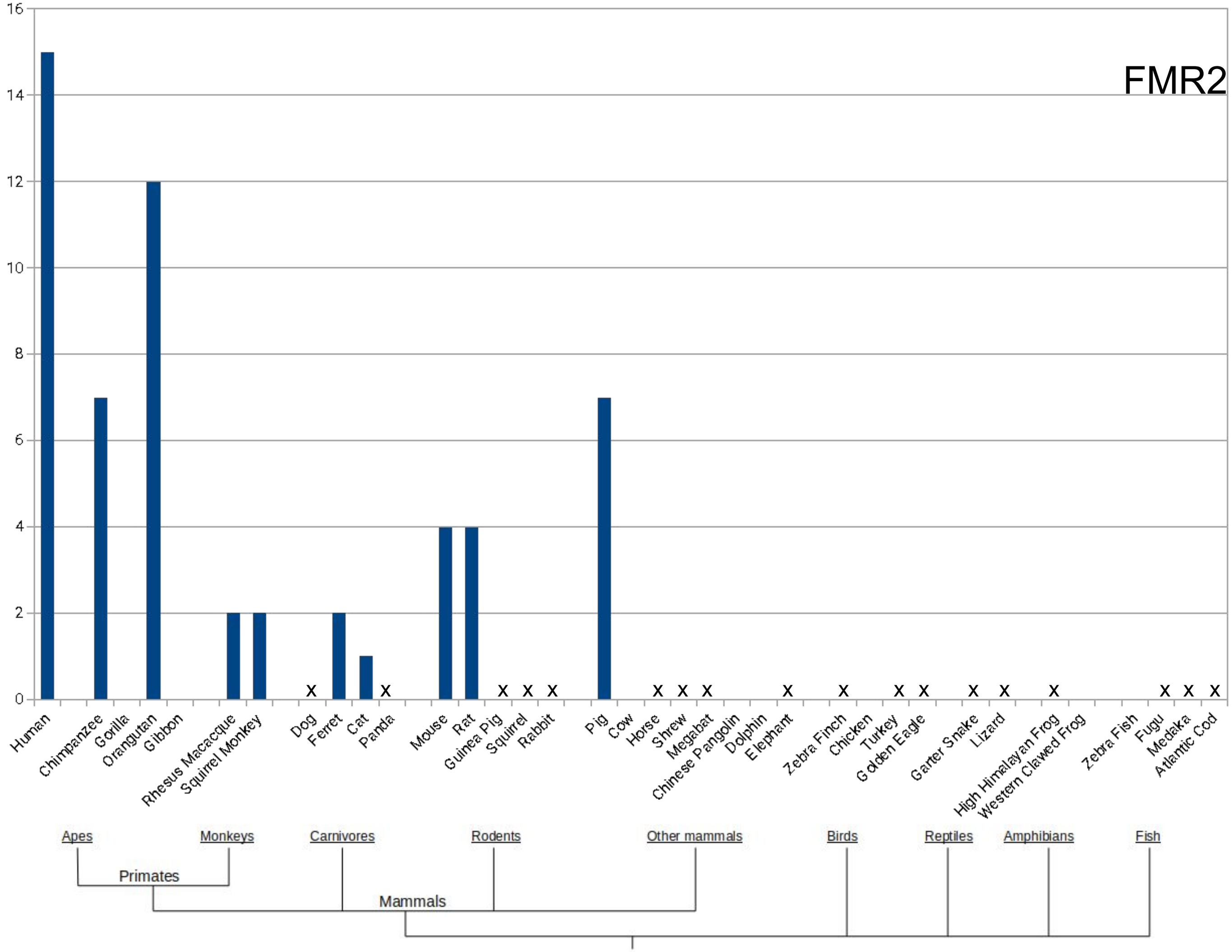

FMR2
0
2
4
6
8
10
12
14
16
Human
Chimpanzee
Gorilla
Orangutan
Gibbon
Rhesus Macacque
Squirrel Monkey
Dog
Ferret
Cat
Panda
Mouse
Rat
Guinea Pig
Squirrel
Rabbit
Pig
Cow
Horse
Shrew
Megabat
Chinese Pangolin
Dolphin
Elephant
Zebra Finch
Chicken
Turkey
Golden Eagle
Garter Snake
Lizard
High Himalayan Frog
Western Clawed Frog
Zebra Fish
Fugu
Medaka
Atlantic Cod
Apes
Monkeys
Carnivores
Rodents
Other mammals
Birds
Reptiles
Amphibians
Fish
Primates
Mammals

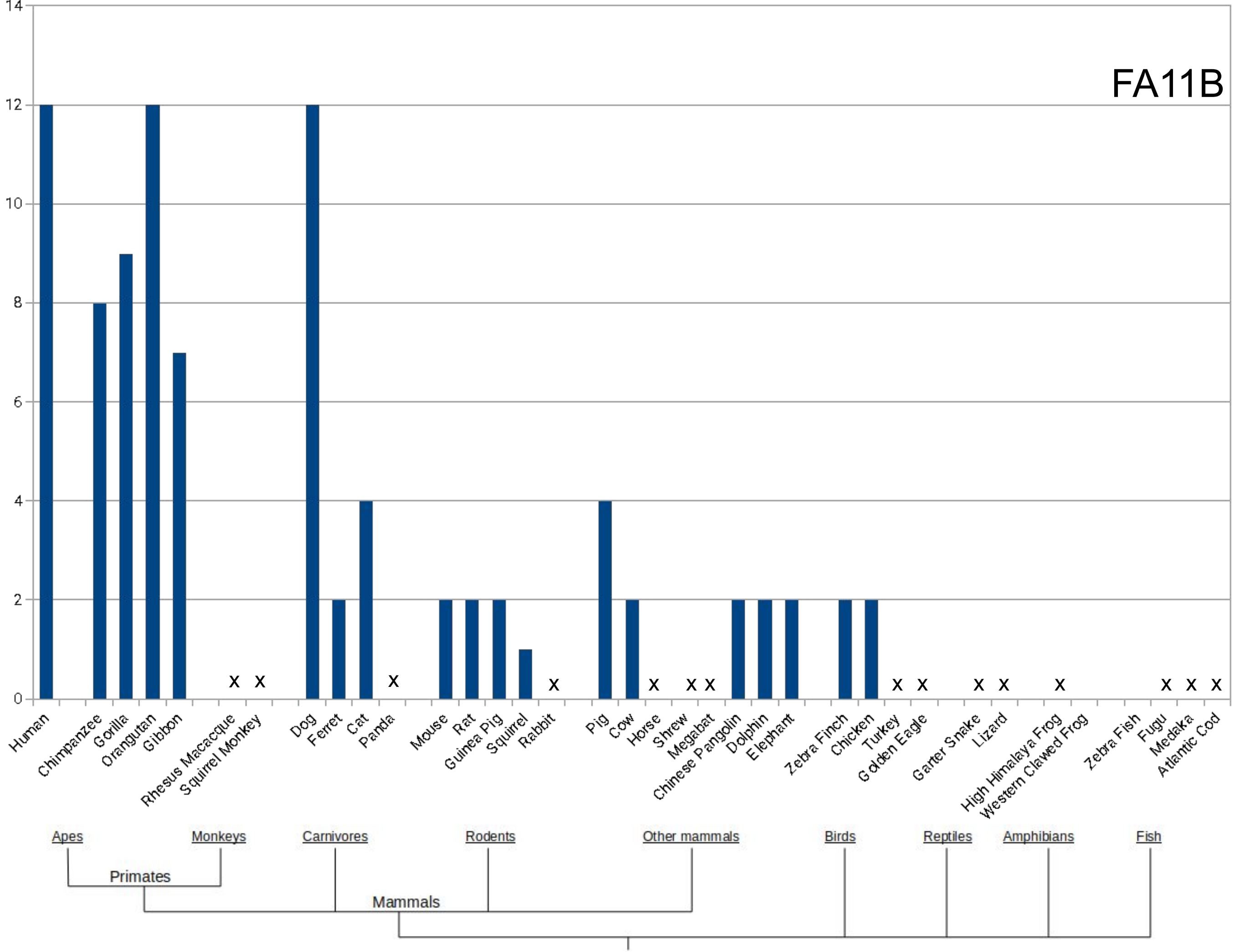

FA11B
14
12
10
8
6
4
2
0
Human
Chimpanzee
Gorilla
Orangutan
Gibbon
Rhesus Macacque
Squirrel Monkey
Dog
Ferret
Cat
Panda
Mouse
Rat
Guinea Pig
Squirrel
Rabbit
Pig
Cow
Horse
Shrew
Megabat
Chinese Pangolin
Dolphin
Elephant
Zebra Finch
Chicken
Turkey
Golden Eagle
Garter Snake
Lizard
High Himalaya Frog
Western Clawed Frog
Zebra Fish
Fugu
Medaka
Atlantic Cod
Apes
Monkeys
Carnivores
Rodents
Other mammals
Birds
Reptiles
Amphibians
Fish
Primates
Mammals
