## Supplementary file 3 for "Expansion of triplet nucleotide repeats in primates and other vertebrates: an evolutionary perspective"

### Length of repeats across vertebrates : OrthoDB approach

### POLYGLUTAMINE REPEAT DISORDERS

### SCA1

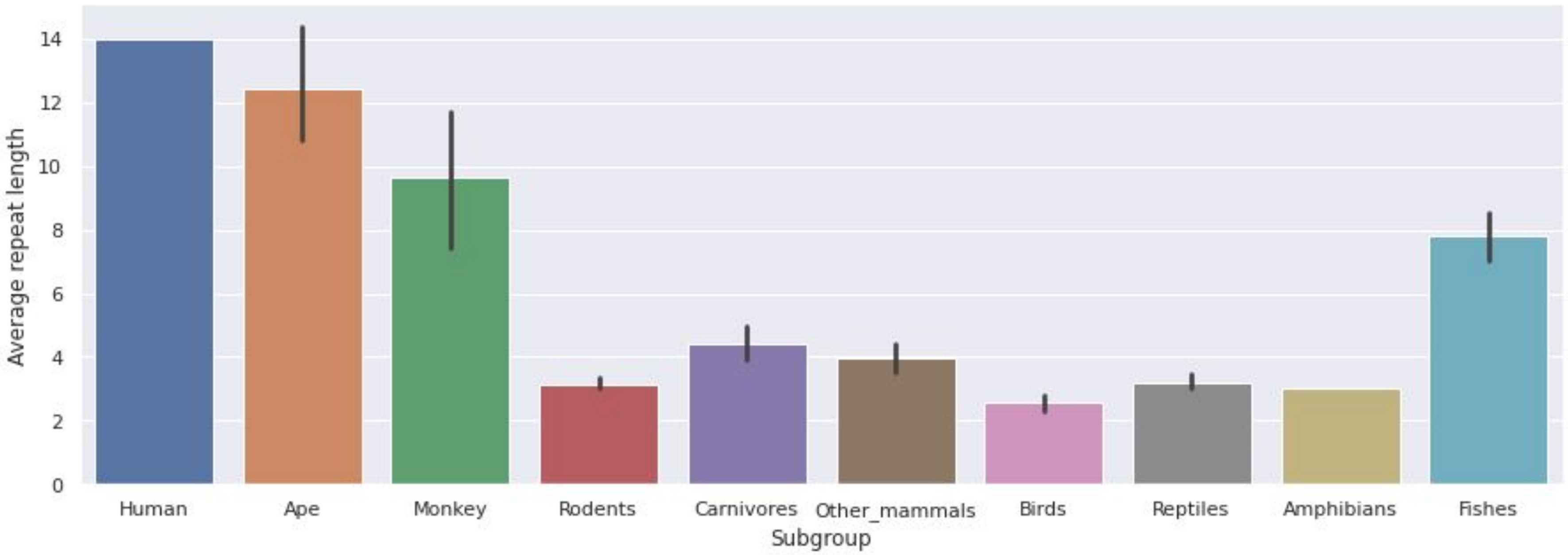
Average repeat length
0
2
4
6
8
10
12
14
Human
Ape
Monkey
Rodents
Carnivores
Other_mammals
Birds
Reptiles
Amphibians
Fishes
Subgroup

### SCA2

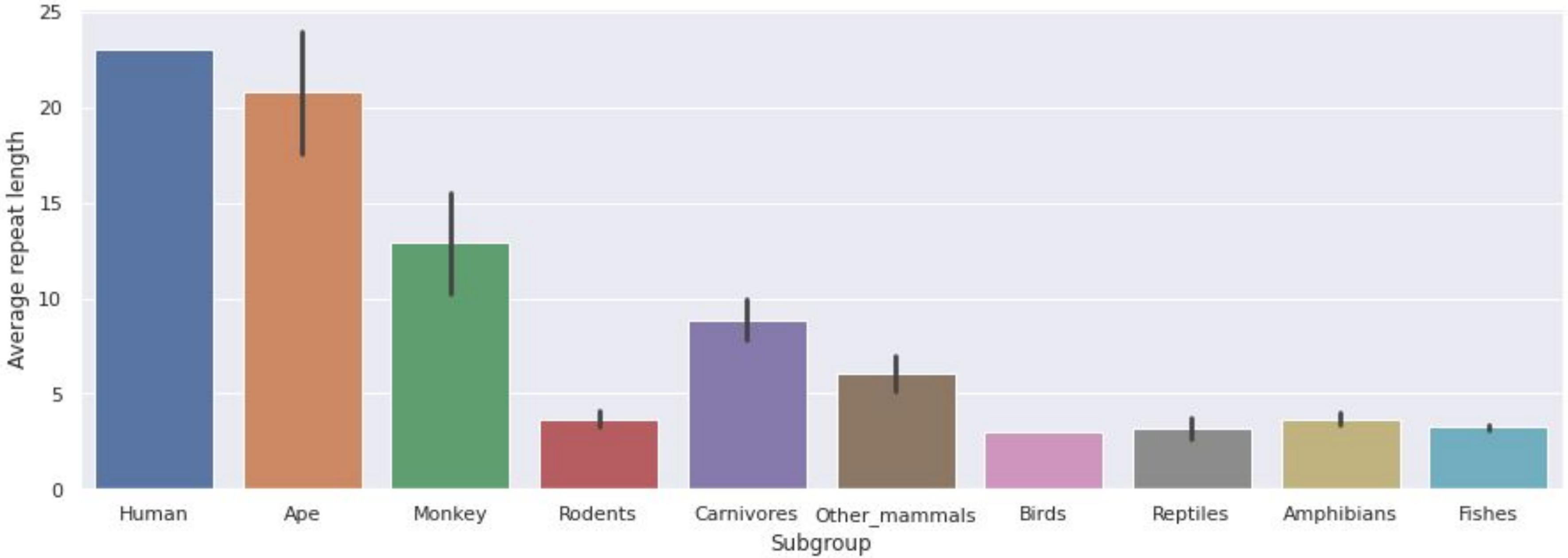

25
20
15
10
5
0
Average repeat length
Human
Ape
Monkey
Rodents
Carnivores
Other_mammals
Birds
Reptiles
Amphibians
Fishes
Subgroup

### SCA3

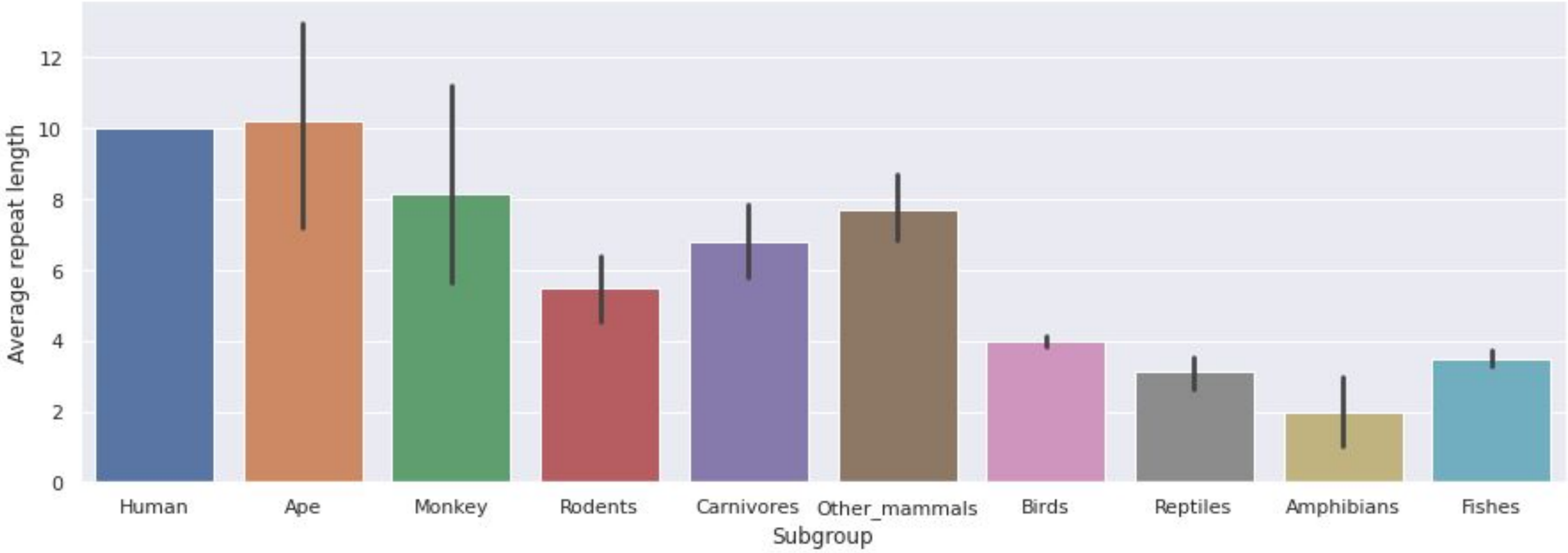

Average repeat length
12
10
8
6
4
2
0
Human
Ape
Monkey
Rodents
Carnivores
Other_mammals
Birds
Reptiles
Amphibians
Fishes
Subgroup

### SCA6

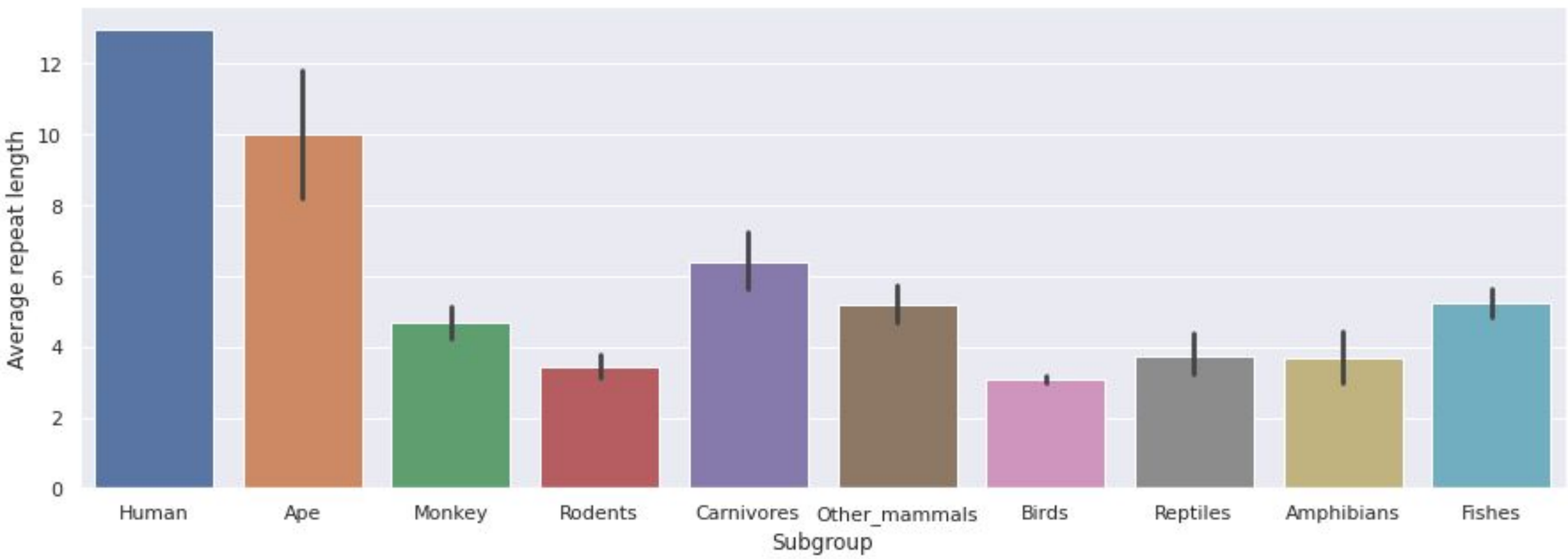

Average repeat length
0
2
4
6
8
10
12
Human
Ape
Monkey
Rodents
Carnivores
Other_mammals
Birds
Reptiles
Amphibians
Fishes
Subgroup

### SCA7

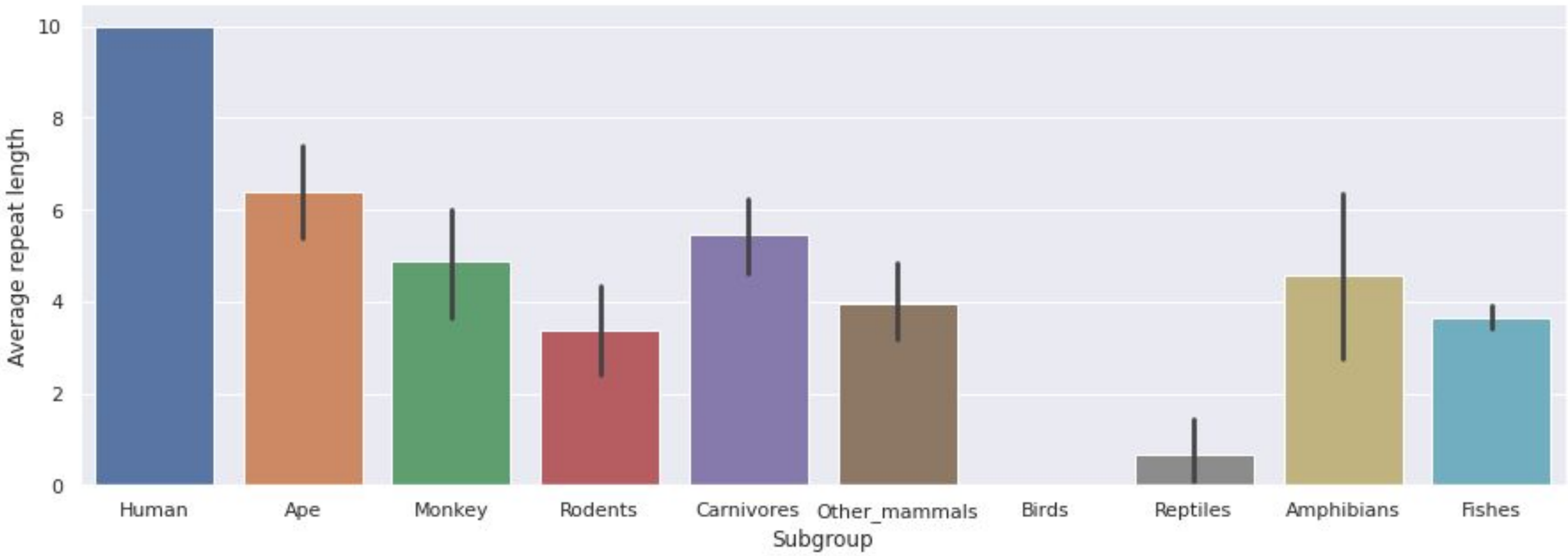

10
8
6
4
2
0
Average repeat length
Human
Ape
Monkey
Rodents
Carnivores
Other_mammals
Birds
Reptiles
Amphibians
Fishes
Subgroup

### SCA17

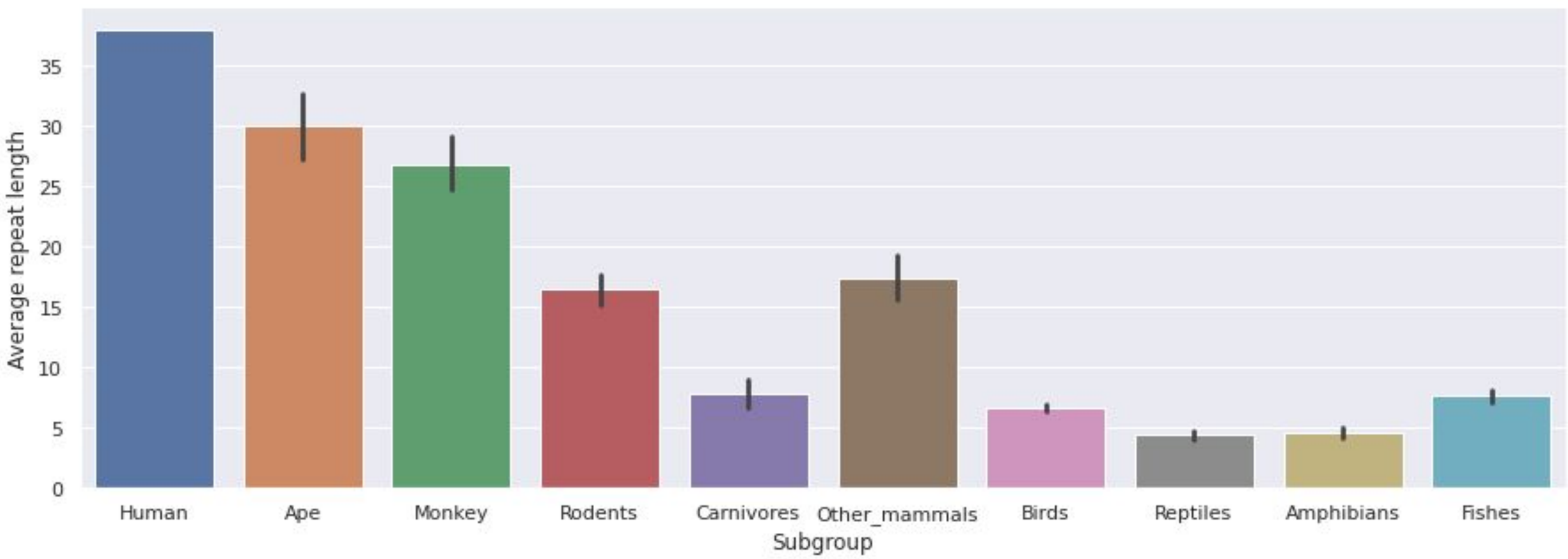
35
30
25
20
15
10
5
0
Average repeat length
Human
Ape
Monkey
Rodents
Carnivores
Other_mammals
Birds
Reptiles
Amphibians
Fishes
Subgroup

### SBMA

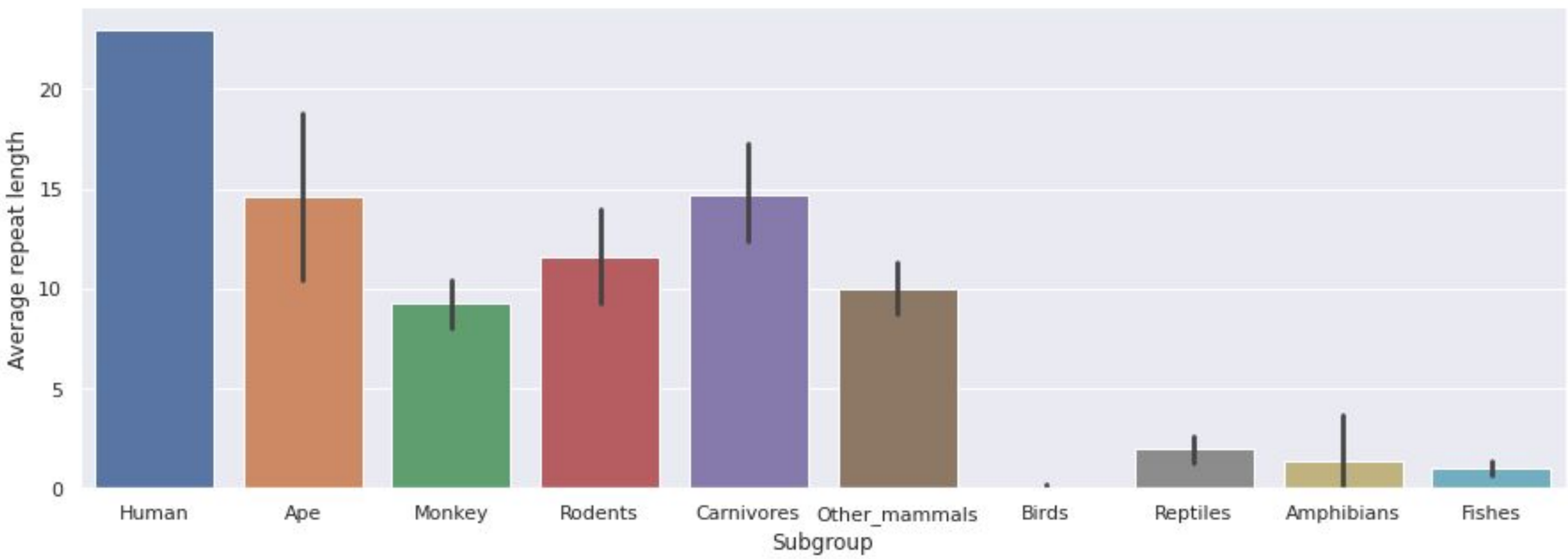

Average repeat length
20
15
10
5
0
Human
Ape
Monkey
Rodents
Carnivores
Other_mammals
Birds
Reptiles
Amphibians
Fishes
Subgroup

# HD

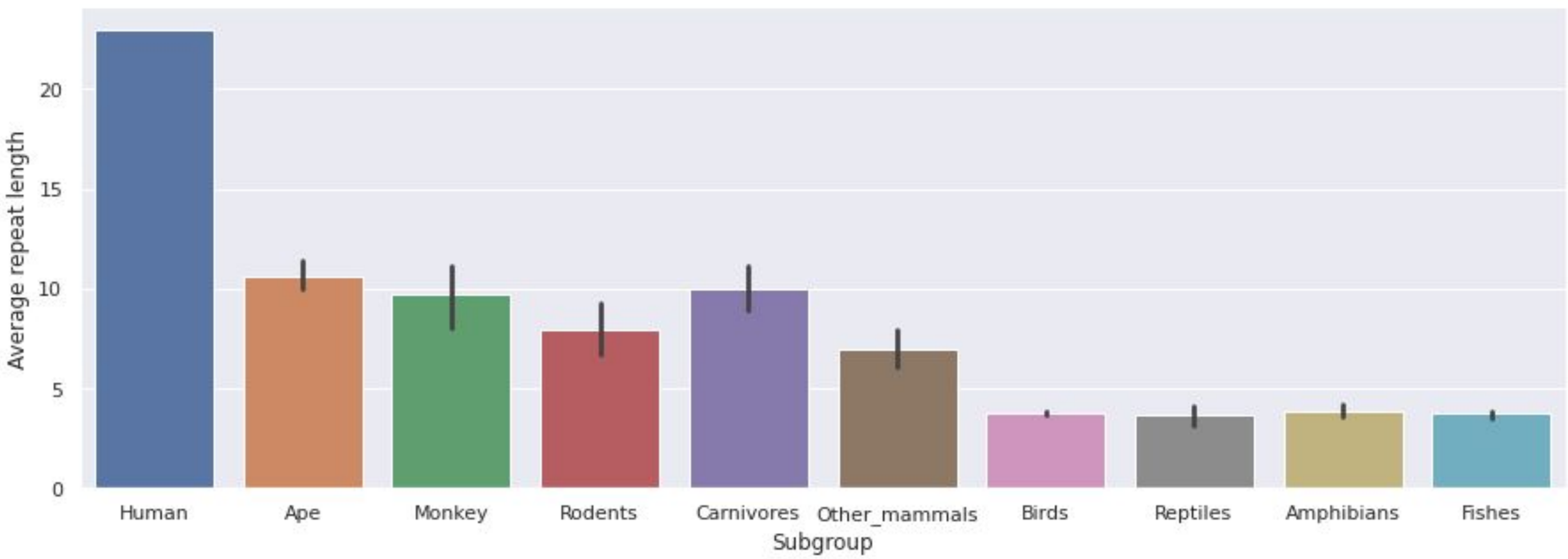
Average repeat length
0
5
10
15
20
Human
Ape
Monkey
Rodents
Carnivores
Other_mammals
Birds
Reptiles
Amphibians
Fishes
Subgroup

### DRPLA

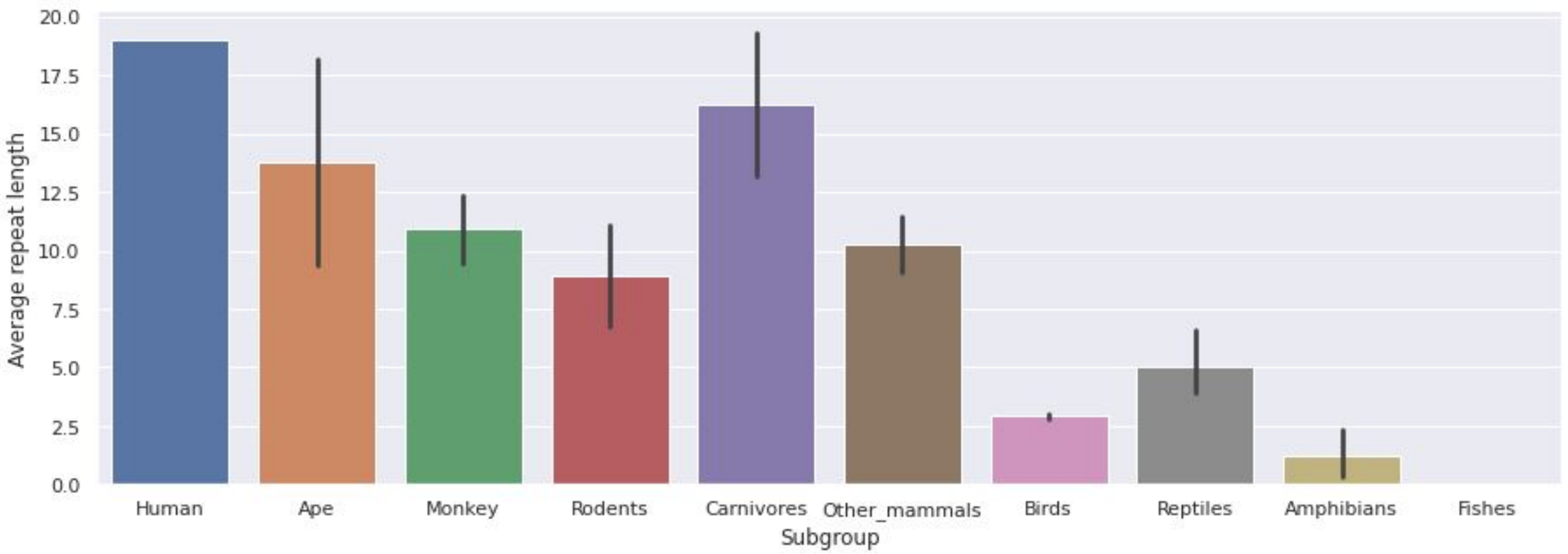

20.0
17.5
15.0
12.5
10.0
7.5
5.0
2.5
0.0
Average repeat length
Human
Ape
Monkey
Rodents
Carnivores
Other_mammals
Birds
Reptiles
Amphibians
Fishes
Subgroup

### KCCN3

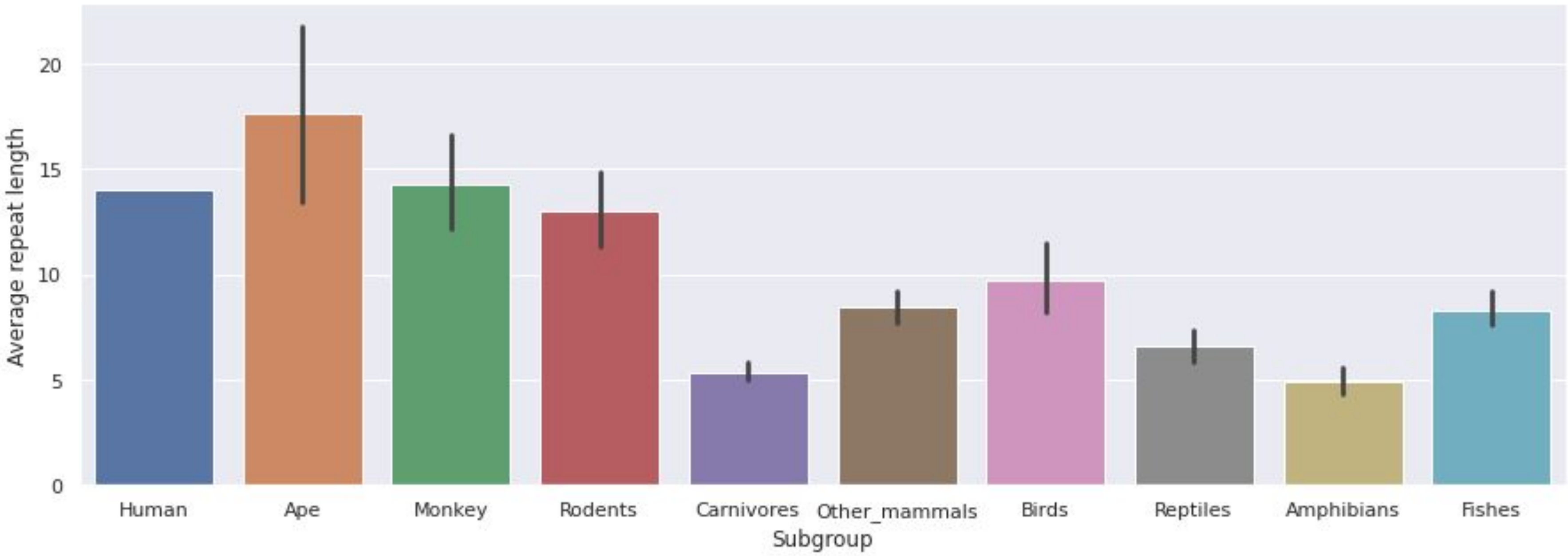

Average repeat length
20
15
10
5
0
Human
Ape
Monkey
Rodents
Carnivores
Other_mammals
Birds
Reptiles
Amphibians
Fishes
Subgroup

### AIB-1

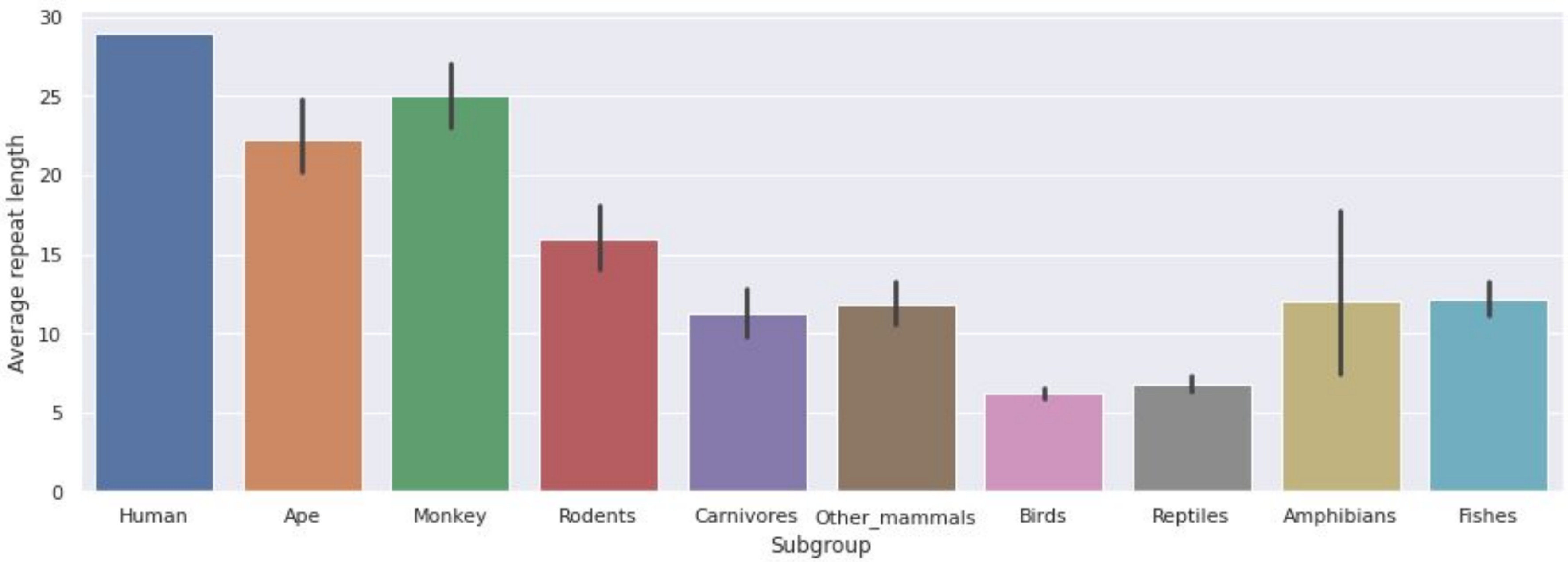

30
25
20
15
10
5
0
Average repeat length
Human
Ape
Monkey
Rodents
Carnivores
Other_mammals
Birds
Reptiles
Amphibians
Fishes
Subgroup

### POLYALANINE REPEAT DISORDERS

### HOXD13

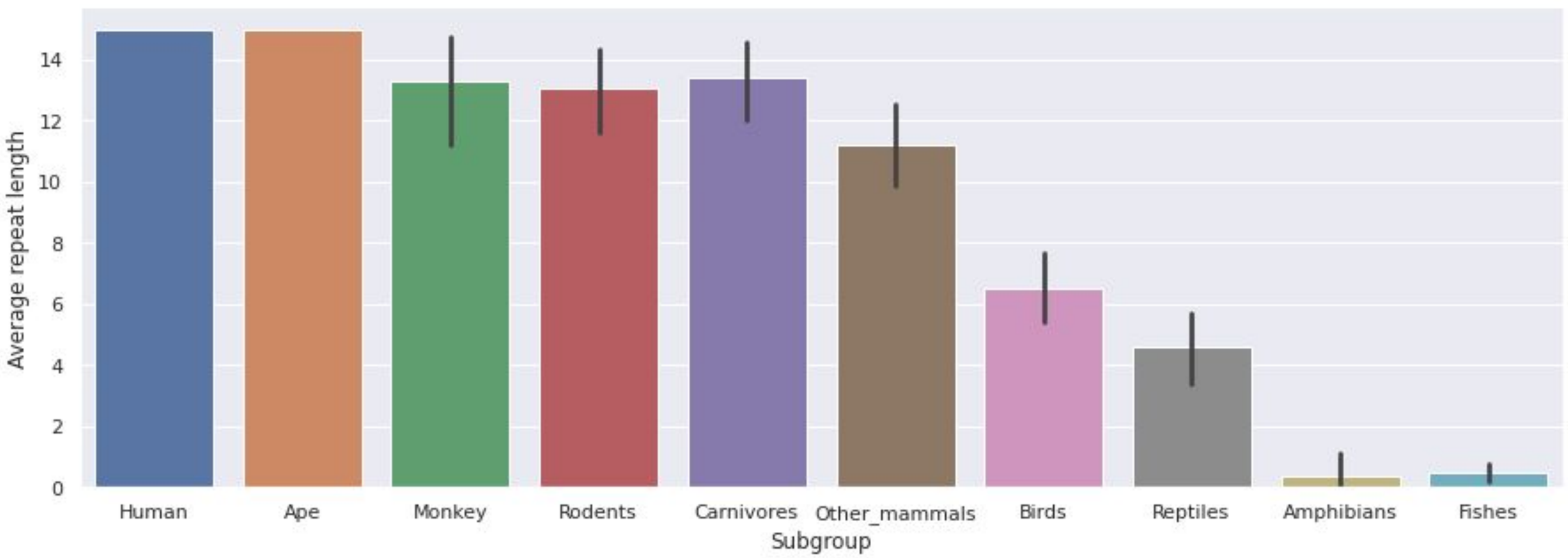

Average repeat length
0
2
4
6
8
10
12
14
Human
Ape
Monkey
Rodents
Carnivores
Other_mammals
Birds
Reptiles
Amphibians
Fishes
Subgroup

**CBFA1**

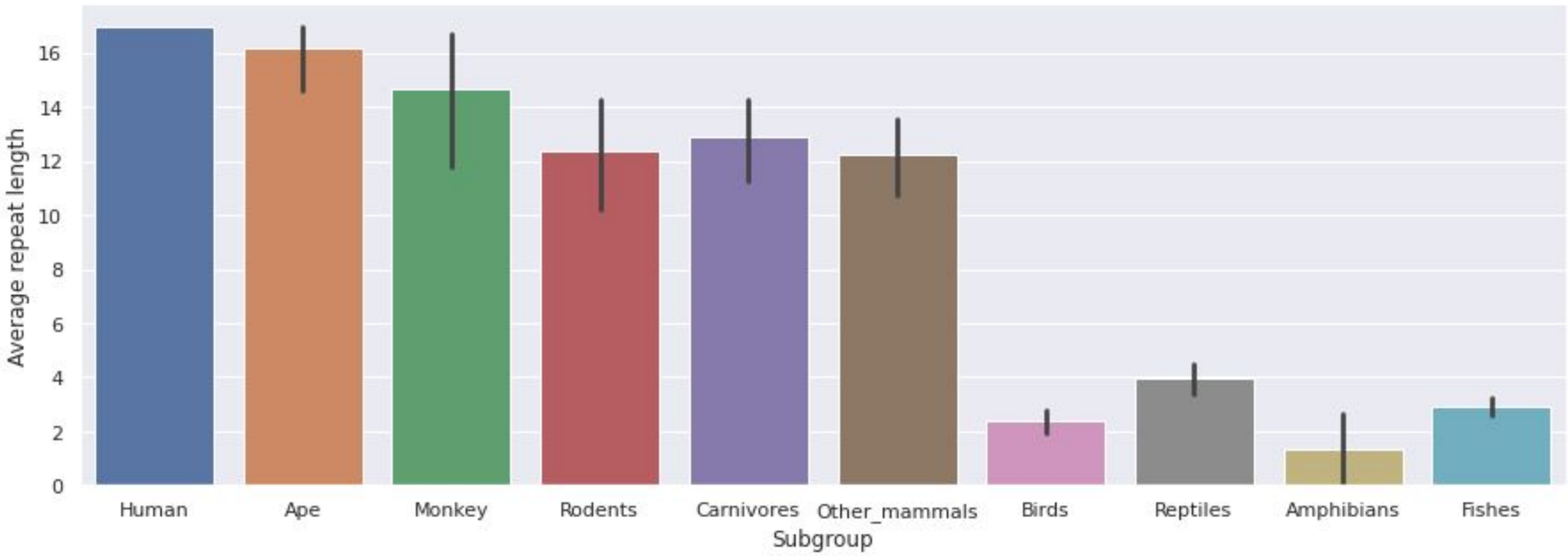

Average repeat length
0
2
4
6
8
10
12
14
16
Human
Ape
Monkey
Rodents
Carnivores
Other_mammals
Birds
Reptiles
Amphibians
Fishes
Subgroup

### ZIC-2

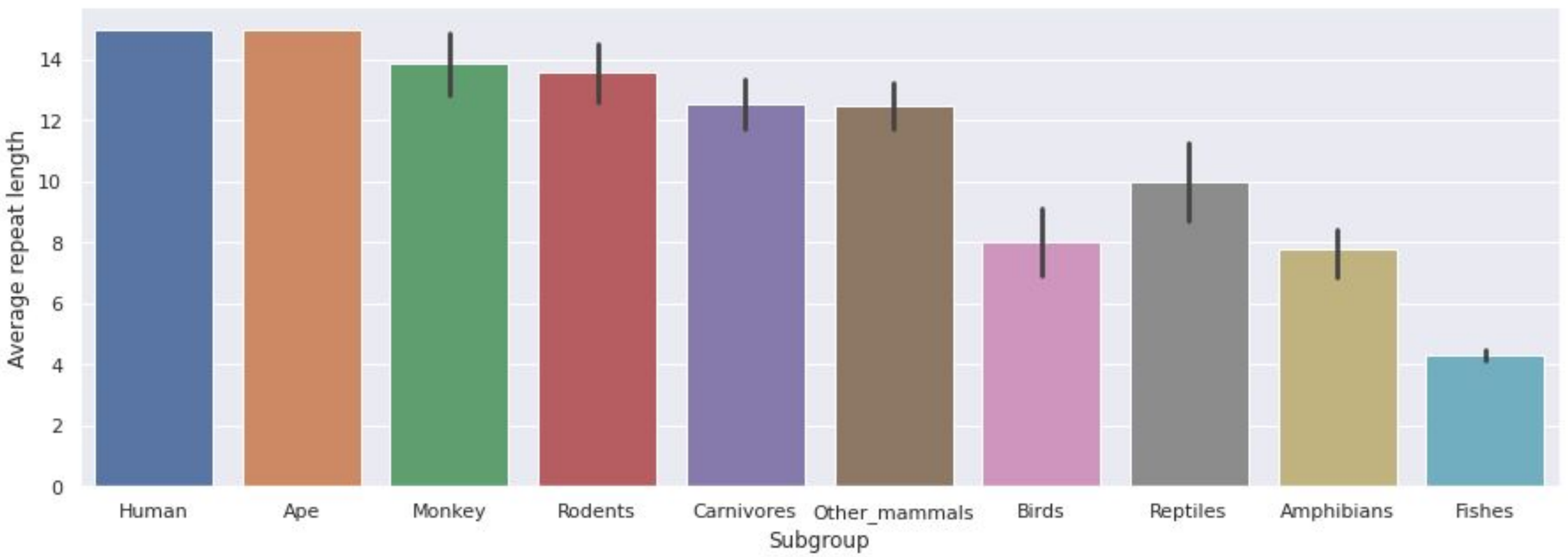
Average repeat length
0
2
4
6
8
10
12
14
Human
Ape
Monkey
Rodents
Carnivores
Other_mammals
Birds
Reptiles
Amphibians
Fishes
Subgroup

### HOXA13

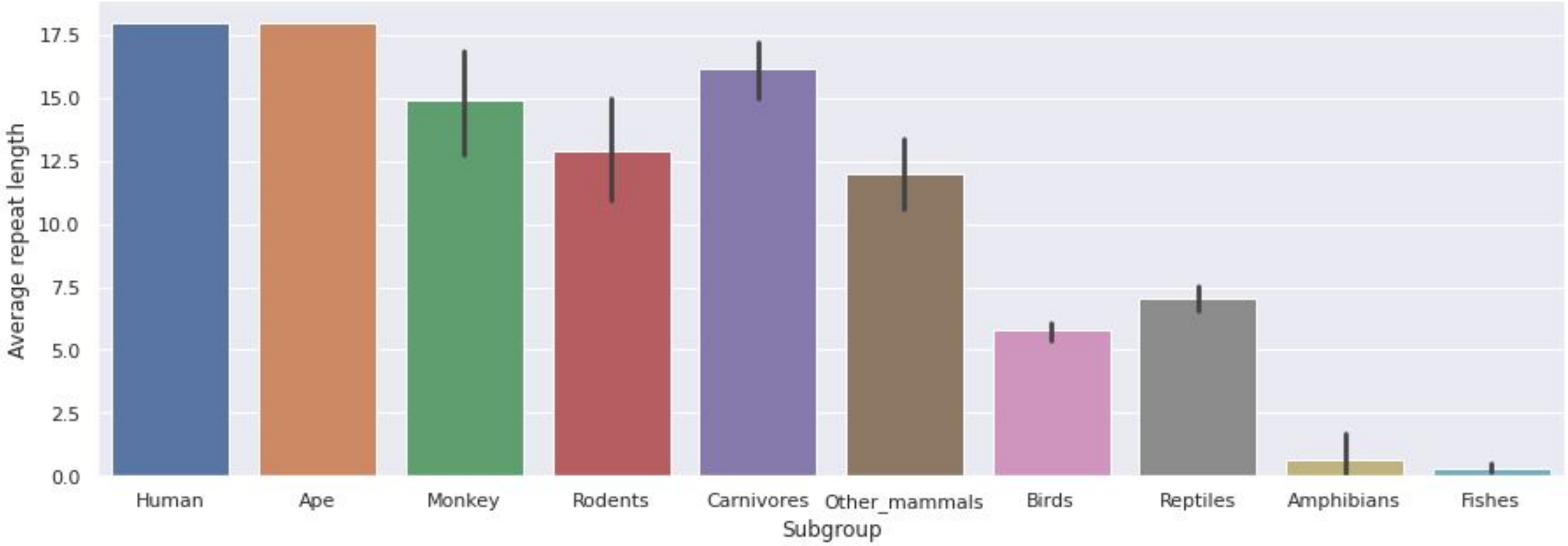

17.5
15.0
12.5
10.0
7.5
5.0
2.5
0.0
Average repeat length
Human
Ape
Monkey
Rodents
Carnivores
Other_mammals
Birds
Reptiles
Amphibians
Fishes
Subgroup

### FOXL2

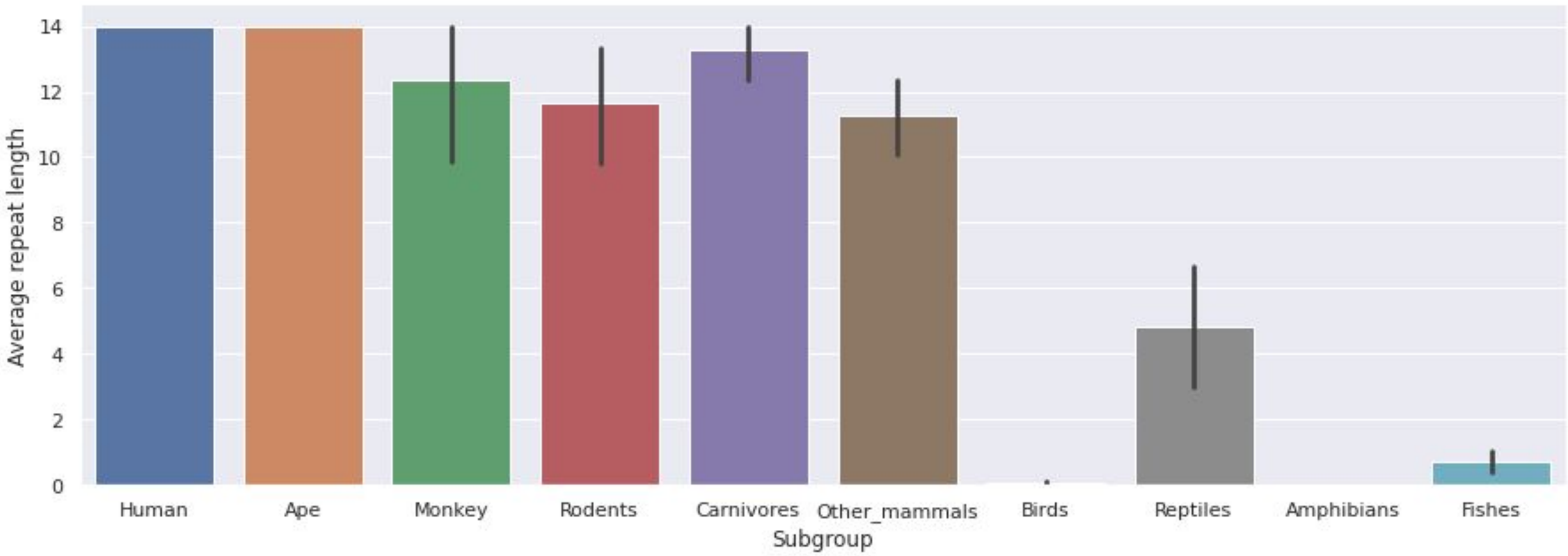

Average repeat length
14
12
10
8
6
4
2
0
Human
Ape
Monkey
Rodents
Carnivores
Other_mammals
Birds
Reptiles
Amphibians
Fishes
Subgroup

### ARX

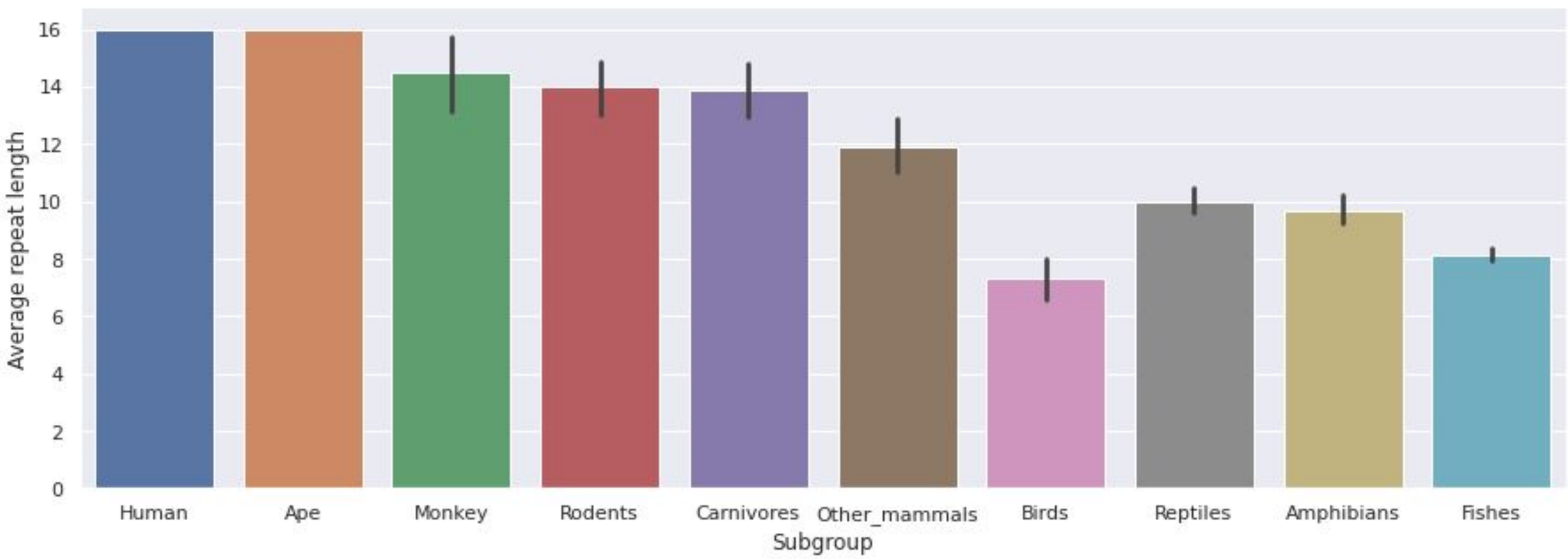

Average repeat length
0
2
4
6
8
10
12
14
16
Human
Ape
Monkey
Rodents
Carnivores
Other_mammals
Birds
Reptiles
Amphibians
Fishes
Subgroup

### POLYASPARTATE REPEAT DISORDER

### COMP

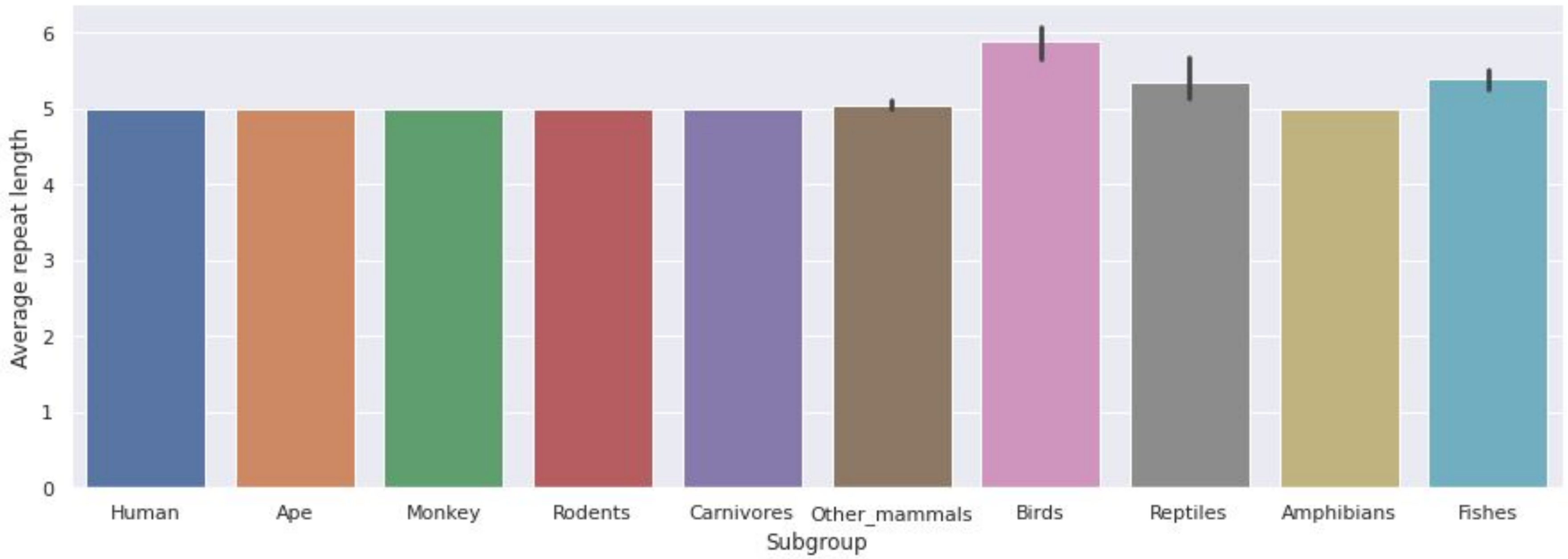
Average repeat length
Subgroup
0
1
2
3
4
5
6
Human
Ape
Monkey
Rodents
Carnivores
Other_mammals
Birds
Reptiles
Amphibians
Fishes
