## Supplementary file 4 for "Expansion of triplet nucleotide repeats in primates and other vertebrates: an evolutionary perspective"

### Trinucleotide repeat disorders : length of repeats across vertebrates

### POLYGLUTAMINE REPEAT DISORDERS - CEREBELLAR ATAXIAS

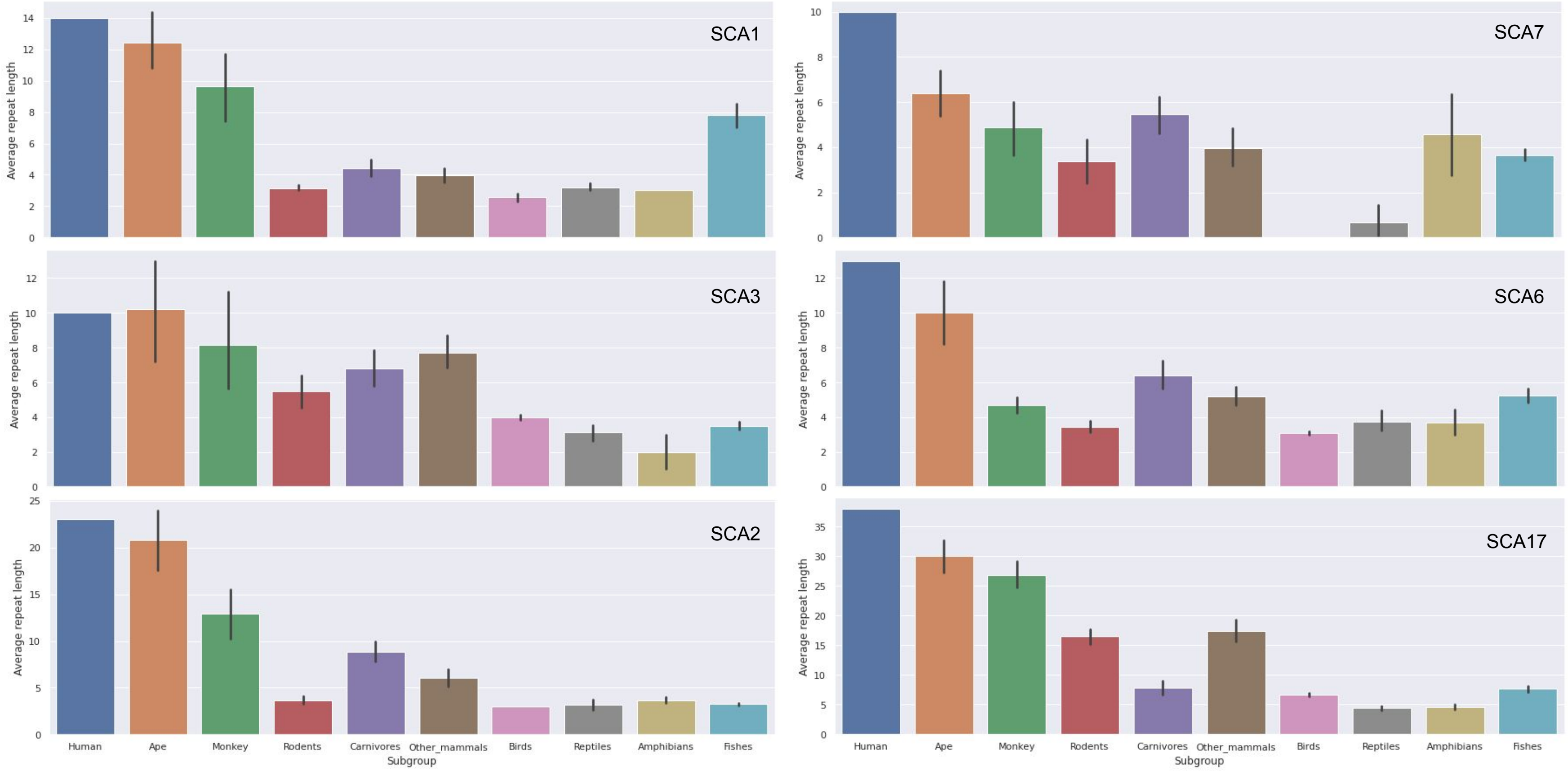

### POLYGLUTAMINE REPEAT DISORDERS - OTHERS

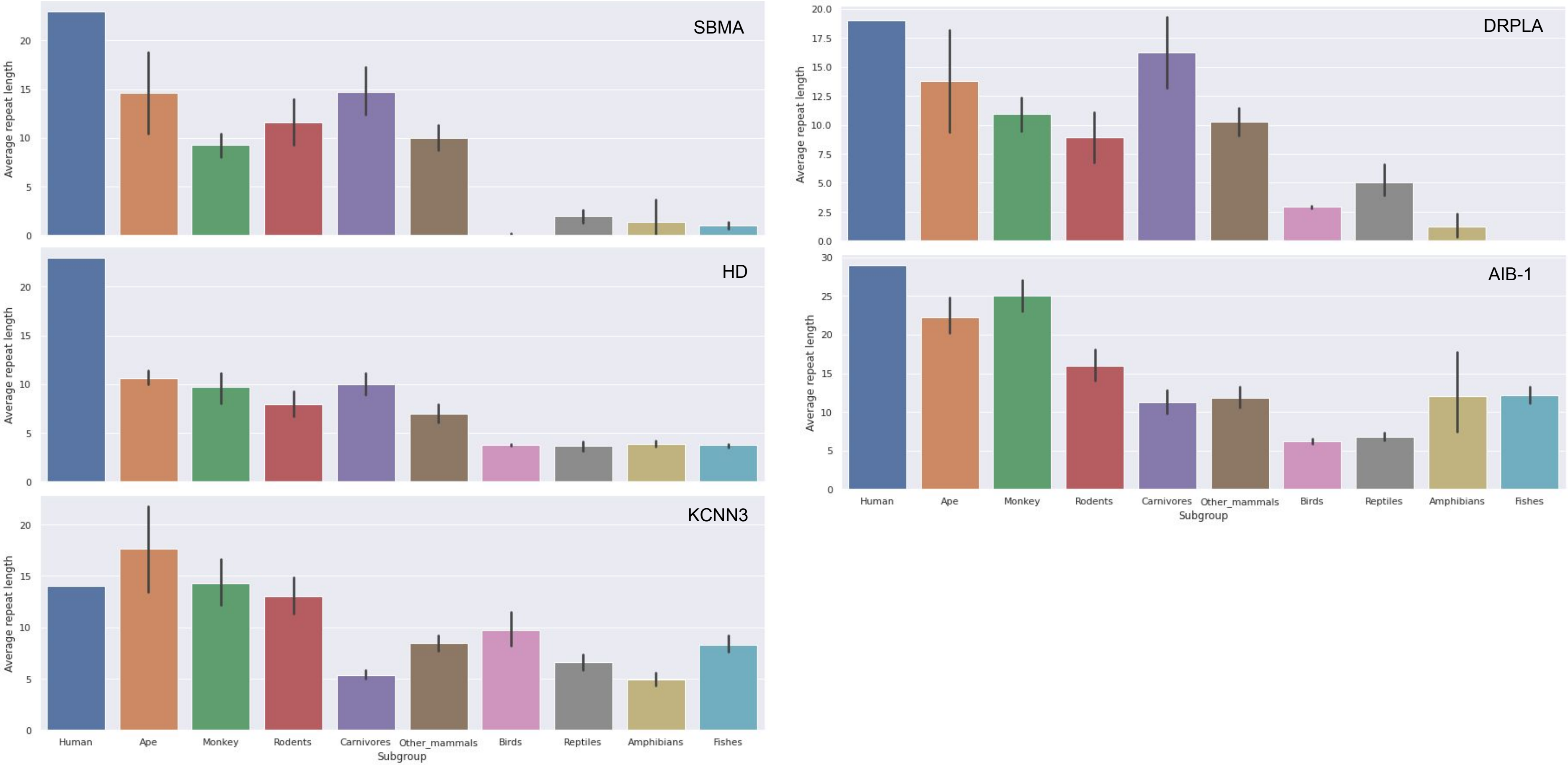

### POLYALANINE REPEAT DISORDERS

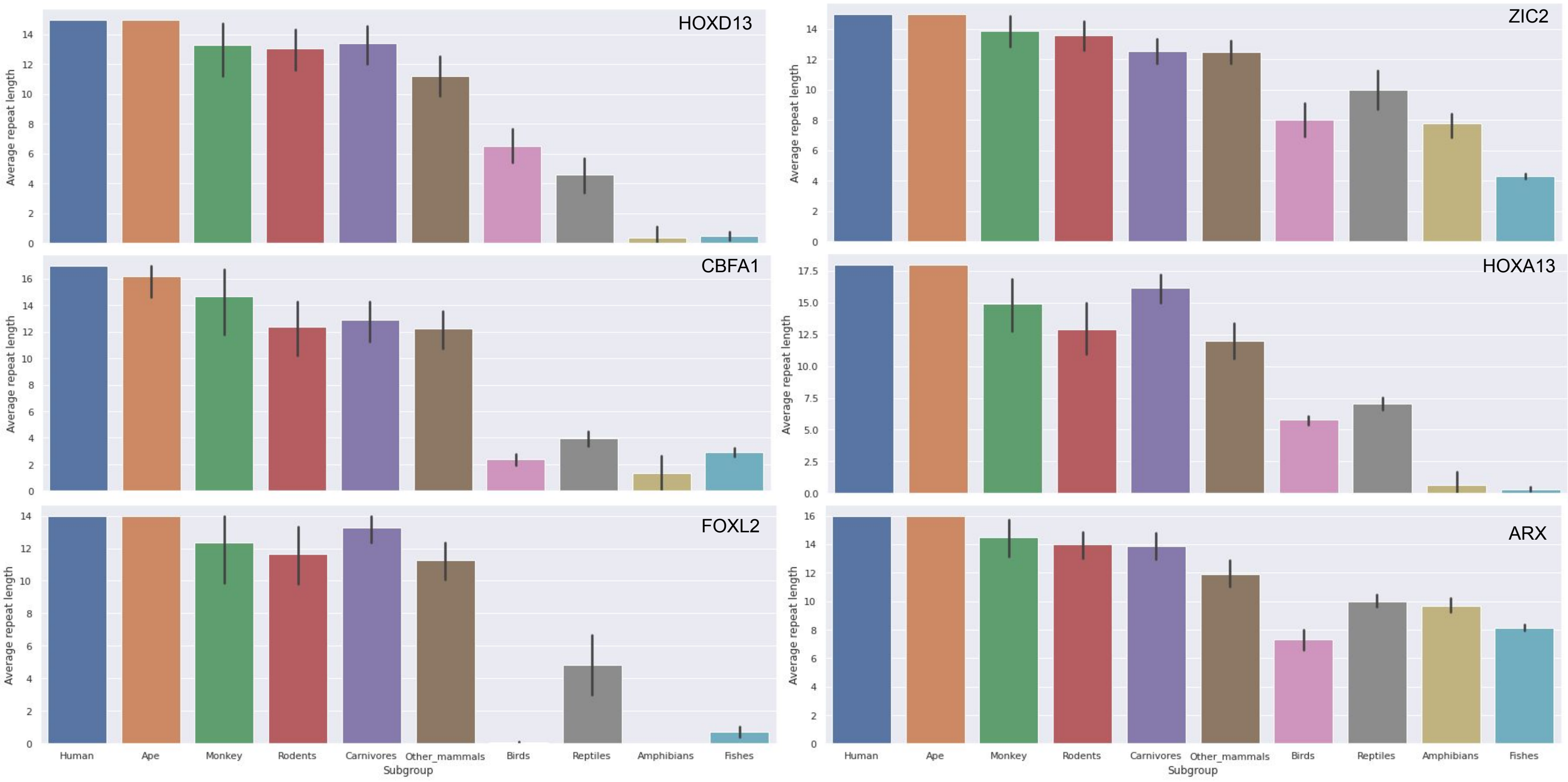

### POLYASPARTATE REPEAT DISORDER

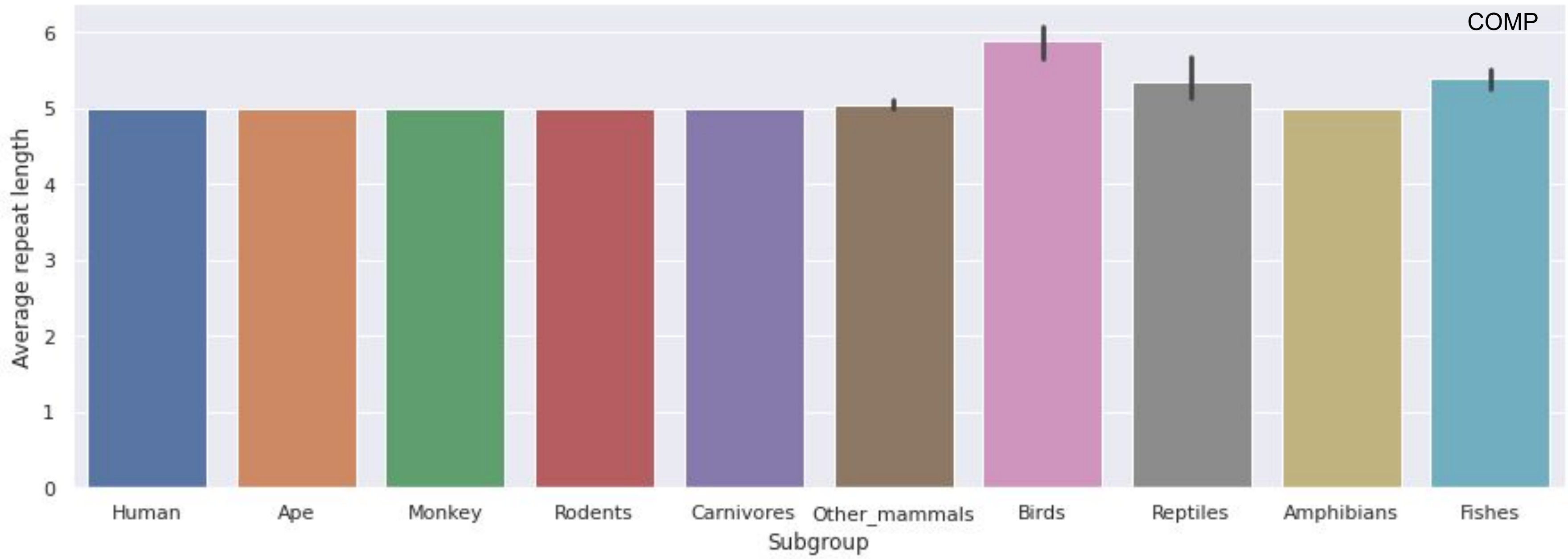
